# Microbiota promote enhanced CD39 expression in γδ intraepithelial lymphocytes through the activation of TCR and IL-15 signaling

**DOI:** 10.1101/2025.03.22.644616

**Authors:** Sara Alonso, Harsimran Kaur, Luo Jia, Mai-Uyen Nguyen, Alyssa Laguerta, Andrew Fong, Neema Skariah, Rafael J. Argüello, Michael P. Verzi, Mahima Swamy, Ken S. Lau, Karen L. Edelblum

## Abstract

Intraepithelial lymphocytes expressing the γδ T cell receptor (γδ IEL) provide continuous surveillance of the intestinal epithelium. We report that mice harboring a microbiota-specific hyperproliferative γδ IEL (γδ^HYP^) phenotype also upregulate the expression of the ectonucleotidase CD39, a marker of regulatory γδ T cells. Enhanced TCR and IL-15 signaling correlates with a progression from a naïve-like CD39^neg^ γδ IEL to a more mature, tissue-adapted CD39^hi^ IEL population. We found that TCRγδ activation drives CD122-mediated CD39 upregulation on γδ^HYP^ IELs and increased mucosal IL-15 further amplifies CD39 expression in these cells. Further investigation revealed that CD39 induction requires sustained exposure to the γδ^HYP^-associated microbiota. Moreover, CD39^hi^ γδ IELs exhibit a reduced capacity to produce pro-inflammatory cytokine, which may explain the lack of histopathology in γδ^HYP^ mice. Overall, our study identifies a previously unappreciated mechanism by which an altered microbiota amplifies CD39 expression on γδ^HYP^ IELs, leading to the expansion of γδ IELs with regulatory potential.

## INTRODUCTION

Intraepithelial lymphocytes (IEL) are located within the intestinal epithelium and serve as a first line of defense against infection, injury, and inflammation^1, 2^. IELs are broadly divided into two main subsets: induced IELs (CD4 TCRαβ, CD8αβ TCRαβ or CD4 CD8αα), which are conventional T cells that home to the gut from the periphery, and unconventional, natural IELs (CD8αα TCRαβ or TCRγδ) that can be activated in an MHC-independent manner. TCRγδ IELs (γδ IEL) comprise approximately 60% of the murine IEL compartment and dynamically survey the epithelial barrier^3, 4^. Although γδ IELs have cytotoxic potential, these cells maintain an ‘activated yet resting’ phenotype to allow for a rapid response to microbial infection while limiting the likelihood for aberrant activation^2^. The molecular mechanisms that regulate or restrain γδ IEL effector function remain an area of ongoing investigation.

Natural IELs are highly dependent on IL-15 for a variety of cellular functions^5, 6^. This pleiotropic cytokine is produced by intestinal epithelial cells and lamina propria dendritic cells in response to the commensal-mediated activation of pattern recognition receptors (PRR)^7–10^. IL-15 complexes with IL-15 receptor α (IL-15RC), which once trafficked to the plasma membrane, is presented in *trans* to IL-2Rβ (CD122) on IELs^11^. IL-15 signaling contributes to γδ IEL motility^12^, promotes IEL survival through the upregulation of the anti-apoptotic factor Bcl-2^13, 14^, and supports cellular growth and proliferation through increased nutrient and amino acid (AA) uptake, metabolism, and protein production^5^. Thus, it is not surprising that mice deficient in IL-15, IL-15Rα, or IL-2Rβ fail to maintain an intact γδ IEL compartment^13, 15–17^.

γδ IELs can develop in the absence of commensal bacteria^18, 19^; however, several microbial-derived factors contribute to the maintenance of these sentinel lymphocytes. In addition to inducing PRR-dependent IL-15 production^7–10^, commensal bacteria generate tryptophan metabolites, such as indoles, to activate the aryl hydrocarbon receptor (AhR) expressed by IELs^20^. Defective AhR signaling leads to impaired maintenance of the IEL compartment^20, 21^. Recently, microbial-derived indoles were shown to be recognized by a polyspecific TCRγδ^22^, yet how γδ IELs recognize and expand in response to commensals requires further study. We previously identified mice in our facility harboring an altered microbiota that was sufficient to drive a hyperproliferative γδ IEL (γδ^HYP^) phenotype^23^. Further, these expanded γδ^HYP^ IELs were highly motile and conferred protection against systemic *Salmonella* infection. Increased IEL number is often associated with intestinal pathology, including celiac disease and microscopic colitis^24, 25^. Despite the substantial increase in IEL number, we were surprised to find that γδ^HYP^ mice show no signs of overt intestinal pathology, indicating that γδ^HYP^ IELs may possess additional regulatory or compensatory mechanisms to prevent tissue damage or inflammation.

CD39 (encoded by *Entpd1*) is a cell surface ectonucleotidase that rapidly hydrolyzes extracellular ATP into ADP or AMP^26^. These molecules are subsequently hydrolyzed by CD73 into adenosine, which upon binding to its cognate receptor(s), can suppress immune cell proliferation and effector cytokine production. Although CD39 expression is typically associated with regulatory T cells and exhausted T cells^27^, CD39 also serves as a marker of regulatory γδ T cells in various tissues^28, 29^. We recently reported that CD39^+^ γδ IELs can suppress activated CD8 T cells in an adenosine-dependent manner, and that these immunoregulatory IELs are decreased prior the onset of chronic ileitis^30^. Despite the observation that IL-15 induces CD39 expression in NK cells^31^, the molecular mechanism that drives CD39 expression in γδ IELs is unclear.

We now report that γδ^HYP^ IELs exhibit an expansion of a CD39^hi^ population that is generated in response to increased TCR and IL-15 signaling. Activation of these pathways correlates with a progression from a naïve-like CD39^neg^ γδ IEL to a more mature, tissue-adapted CD39^hi^ IEL population. Using *in vitro* assays coupled with antibody-mediated blockade *in vivo*, we find that TCRγδ activation drives CD122-mediated upregulation of CD39 on γδ^HYP^ IELs. Further, sustained exposure to the γδ^HYP^-associated microbiota is required for the expansion of CD39^hi^ γδ IELs. We show that enhanced IL-15 signaling also extends the survival of γδ^HYP^ IELs and likely enhances the bioenergetic potential of these cells. Notably, there was an inverse relationship between CD39 expression and pro-inflammatory cytokine production, suggesting that an increase in CD39^hi^ γδ IELs may explain the absence of histopathology in γδ^HYP^ mice. Our study identifies a novel mechanism by which an altered microbiota enhances γδ IEL CD39 expression, thus creating an environment that promotes the generation of immunosuppressive γδ IELs.

## RESULTS

### γδ^HYP^ mice exhibit an expansion of a CD39^hi^ population

We previously reported the identification of a mouse line in our colony harboring a unique microbiota that was both necessary and sufficient to promote substantial increase in γδ IEL number relative to wildtype (WT) mice^23^. Further, vertical transfer of the microbiota by crossing TcrdEGFP (WT) dams with phenotypic interferon alpha receptor (IFNAR)-deficient sires led to the generation of phenotypic F2 WT littermates, subsequently referred to as γδ^HYP^ mice. As expected, γδ^HYP^ mice exhibit a 3-fold increase in γδ IEL number compared to WT controls (Fig. 1A,B). Profiling of the IEL compartment in γδ^HYP^ mice revealed a shift among TCRαβ IELs favoring an expansion of natural CD8αα TCRαβ IELs and reduction in induced CD8αβ TCRαβ IELs (Supplementary Fig. 1A,B). However, the relative proportion of CD4 TCRαβ (CD4), CD4 CD8αα and CD8α^-^ CD8β^-^ TCRγδ^+^ (double negative, DN) IELs were unchanged between the two phenotypes. Given our interest in γδ IELs, we analyzed the Vγ subsets and observed no difference in the frequency of Vγ7, Vγ1, or Vγ4 IELs between phenotypic and non-phenotypic mice (Supplementary Fig. 1C). Together, the increase in γδ IEL number coupled with the similar overall proportion of TCRαβ and TCRγδ IELs suggests an expansion of the IEL compartment in γδ^HYP^ mice with a slight shift towards natural IELs.

**Figure 1.**
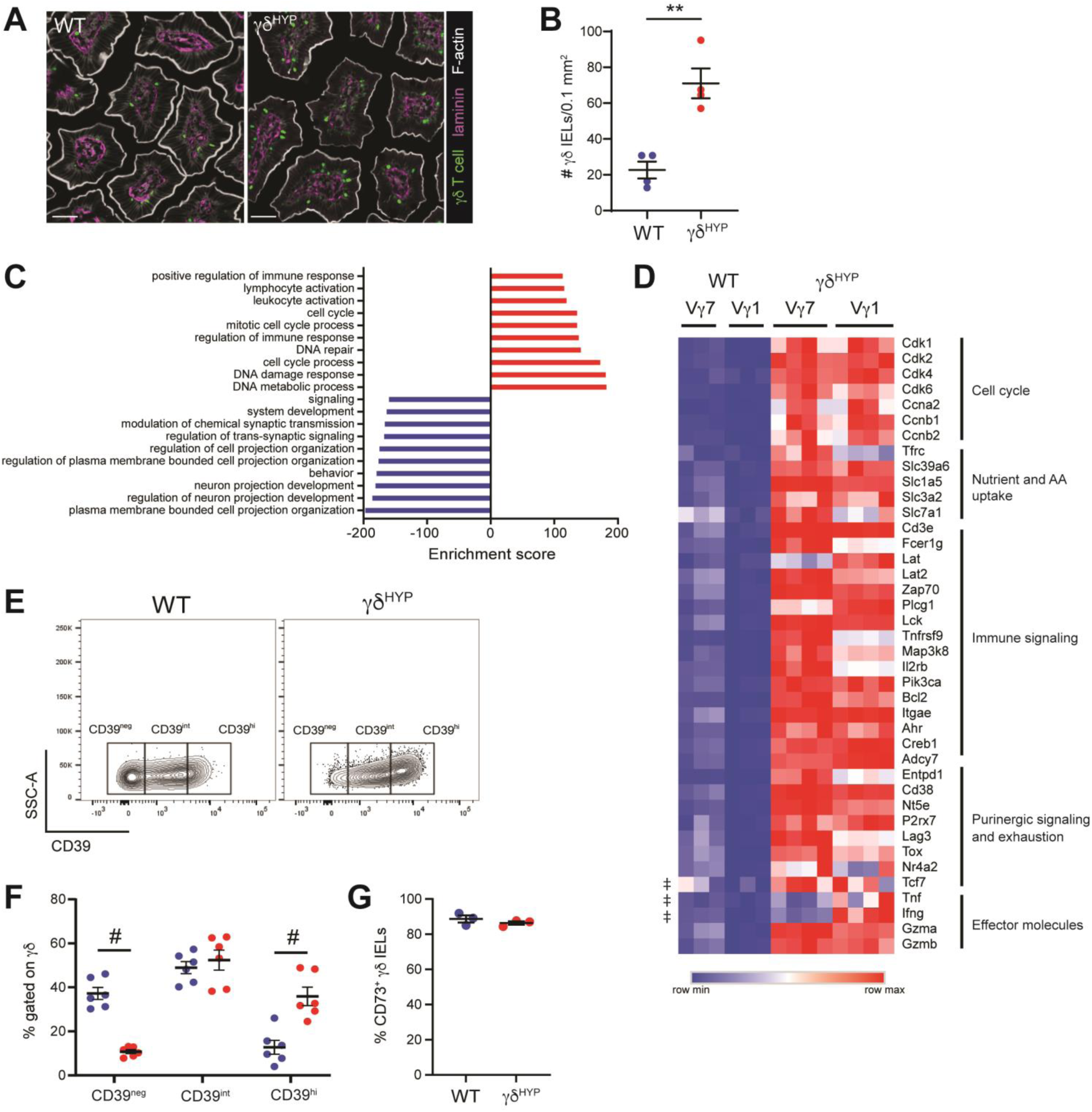
γδ^HYP^ mice exhibit an expansion of a CD39^hi^ population. (A) Immunofluorescence micrographs showing γδ IELs (green) in WT and γδ^HYP^ jejuna. Laminin is shown in magenta and F-actin in white. Scale bar = 30 μm. (B) Morphometric analysis of jejunal γδ IELs in WT and γδ^HYP^ mice. Bulk RNAseq was performed on sorted Vγ7 and Vγ1 IELs isolated from WT and γδ^HYP^ mice. n=3-4. (C) GO terms of biological processes up and downregulated in γδ IELs (combined Vγ7 and Vγ1 IELs) from γδ^HYP^ mice compared to WT (gene sets <1,000 genes). (D) Heatmap of relative gene expression comparing WT vs γδ^HYP^ Vγ7 IELs and WT vs γδ^HYP^ Vγ1 IELs. Genes pertaining to cell cycle, nutrient and AA uptake, immune signaling, purinergic signaling and exhaustion, and effector molecules are shown. (E) Representative flow cytometry plots of CD39 expression in WT and γδ^HYP^ IELs. (F) Frequency of CD39^neg^, CD39^int^, and CD39^hi^ γδ IELs or (G) CD73^+^ γδ IELs in WT and γδ^HYP^ mice. All data shown as mean ± SEM from at least 2 independent experiments. Each data point represents an individual mouse. n=3-6. Statistical analysis: (B,G) unpaired student’s t-test, (C,D) genes shown exhibit ± 1.5 fold change, DESeq2 adj p-value < 0.05, indicates significance only in γδ^HYP^ Vγ1 relative to WT, (F) two-way ANOVA with Tukey’s post hoc test. **P<0.01, #P<0.0001.

Despite a substantial increase in IEL number, the absence of histopathology in γδ^HYP^ mice led us to hypothesize that additional regulatory or compensatory mechanisms may be in place to restrain IEL activation. To interrogate how γδ^HYP^ IELs may be regulated, we assessed the transcriptomic signature of sorted Vγ7 and Vγ1 IELs isolated from WT and γδ^HYP^ mice. Analysis of the combined γδ IEL populations by phenotype, revealed 11,653 differentially expressed genes (DEG) between WT and γδ^HYP^ IELs, with 6,996 genes upregulated in WT and 4,657 upregulated in γδ^HYP^ IELs. Gene ontology (GO) analysis revealed that the top pathways in γδ^HYP^ IELs expressed genes associated with positive activation of the immune system, cell cycle, DNA damage and DNA metabolism, whereas genes related to cellular communication were increased in WT γδ IELs (Fig. 1C). When evaluating the relative expression of select genes, we found that expression patterns were strikingly similar between Vγ subsets regardless of phenotype (Fig. 1D, Supplementary Fig. 2). In line with the GO analysis, both Vγ7 and Vγ1 γδ^HYP^ IELs showed an increase in the expression of genes involved in cell cycle progression, including several cyclins (*Ccna2*, *Ccnb1*, *Ccnb2*) and cyclin-dependent kinases (*Cdk1*, *Cdk2*, *Cdk4*, *Cdk6*). We also observed an upregulation of genes associated with nutrient uptake, such as the iron transporter transferrin (*Tfrc*), the zinc transporter Zip6 (*Slc39a6*), and several amino acid transporters including *Slc1a5*, *Slc3a2*, and *Slc7a1*. These data suggest that γδ^HYP^ IELs may upregulate nutrient transport to support increased proliferation.

The increase in genes involved in immune signaling in γδ^HYP^ IELs led us to further investigate which signaling pathways may be most affected in phenotypic mice. We found that genes associated with the TCR signalosome, such as *Cd3e*, *Lat*, *Zap70*, *Plcg1*, and *Lck* are upregulated in γδ^HYP^ IELs, as are those downstream of TCR activation, including *Tnfrsf9*, *Map3k8*, and *Il2rb* (Fig. 1D). Two negative regulators of TCR signaling, *Lat2* and *Fcer1g*^32^, were also highly expressed, suggesting that γδ^HYP^ IELs may also attenuate activating signals. Several genes involved in purinergic signaling, such as *Entpd1* (CD39), *Cd38*, *Nt5e* (CD73), and *P2rx7*, were also upregulated in γδ^HYP^ IELs. Prior proteomics analyses showed that IELs and exhausted T cells share common markers^32^. Both Vγ7 and Vγ1 γδ^HYP^ IELs exhibit increased expression of the inhibitory receptor *Lag3* and the transcription factors *Tox* and *Nr4a2* relative to WT (Fig. 1D). Given the increase in genes associated with both T cell activation and exhaustion, we next assessed genes related to effector function. We observed minimal changes in *Ifng* and *Tnf* expression, with a slight, but significant increase only in Vγ1 γδ^HYP^ IELs. In contrast, *Gzma* and *Gzmb* were substantially upregulated in both Vγ subsets in γδ^HYP^ IELs compared to WT. Collectively, γδ^HYP^ IELs exhibit an increase in both activating and inhibitory genes yet appear to maintain an effector program.

Based on our recent observation that purinergic signaling contributes to the regulatory potential of γδ IELs^30^, we next investigated whether the increase in *Entpd1* and *Nt5e* translated to the protein level. We observed a marked shift in CD39 expression in γδ^HYP^ IELs toward a CD39^hi^ population accompanied by a nearly complete loss of CD39^neg^ cells (Fig. 1E,F). Natural IELs constitute the majority of CD39-expressing IELs, although the proportion of CD8αα TCRαβ IELs among CD39^pos^ (both intermediate and high expressors) and CD39^hi^ populations was slightly higher in γδ^HYP^ mice (Supplementary Fig. 3A-C). CD8αβ IELs were affected to a lesser extent (Supplementary Fig. 3D). The frequency of CD39^hi^ CD4 and CD39^hi^ CD4 CD8αα IELs were increased; however, CD39 expression was only slightly elevated in DN γδ IELs (Supplementary Fig. 3E-G). From these data, we conclude that the most robust upregulation of CD39 occurs among natural IELs in γδ^HYP^ mice. Evaluation of CD39 among Vγ subsets showed that the vast majority of CD39-expressing γδ IELs, including those that were CD39^hi^, expressed the Vγ7 TCR (Supplementary Fig. 4A-D). While CD39 expression was also enhanced in Vγ1 IELs, this upregulation was not observed in Vγ4 IELs (Supplementary Fig. 4E). In contrast, CD73 protein expression remained high among γδ IELs of either phenotype (Fig. 1G).

### CD39 expression positively correlates with γδ IEL maturation from a naïve-like state to tissue-adapted, immunologically quiescent phenotype

We next investigated whether cellular processes or functional changes are associated with γδ IEL CD39 expression and the extent to which these are driven by the γδ^HYP^ phenotype. To this end, we performed cellular indexing of transcriptomes and epitopes (CITE)-seq to assess transcriptomic changes based on CD39 expression. CD3^+^ IELs were sorted from WT and γδ^HYP^ mice and unsupervised clustering revealed 7 distinct clusters (C) (Fig. 2A). Based on expression of *Cd4*, *Cd8a and Cd8b*, C1 and C6 represent induced CD4 and CD8αβ IELs, respectively. Clusters 2 to 5 consisted of natural IEL populations, as determined by *Cd8a*^pos^ *Cd8b*^neg^ (C2, C3) and *Trdc* (C4, C5) expression. Consistent with previous reports^33^, there was no clear segregation between natural TCRαβ and γδ IELs, rather, two distinct natural IEL clusters formed based on *Tcf7* expression (Fig. 2A,B). Based on this, C2 and C5 were designated as CD8αα TCRαβ *Tcf7*^pos^ and CD8αα TCRγδ *Tcf7*^pos^, respectively, while clusters C3 and C4 were denoted as CD8αα TCRαβ *Tcf7*^neg^ and CD8αα TCRγδ *Tcf7*^neg^. Lastly, C7 contained a separate cluster of proliferating IELs with high *Ki67* expression and a sparse cluster of empty droplets were considered artifact (Fig. 2A). As expected, CD39 antibody labeling overlapped with *Entpd1* expression and revealed a clear expansion of CD39^pos^ cells, particularly among natural IELs (C2-C5), in γδ^HYP^ mice (Fig. 2C,D). Strikingly, CD39 was the highest in *Tcf7*^neg^ natural IELs (C3, C4), suggesting an inverse relationship between CD39 and TCF1 expression.

**Figure 2.**
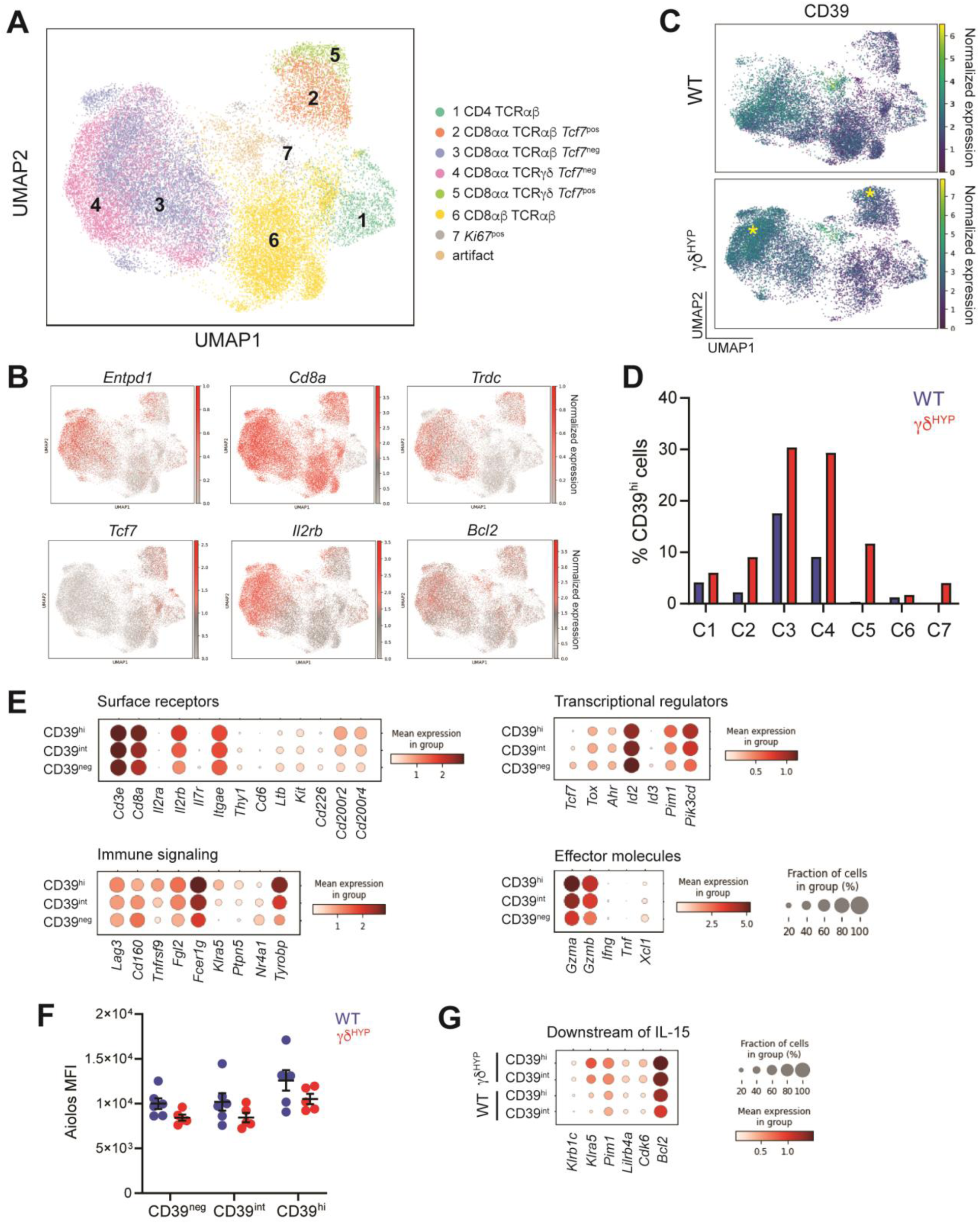
CD39 expression positively correlates with the transition from a naïve-like state to mature, tissue-adapted γδ IELs. CITE-seq was performed on sorted CD3^+^ IELs isolated from WT and γδ^HYP^ mice. (A) scRNA-seq and unsupervised clustering with UMAP analysis representing the distribution of CD3^+^ IELs. (n=3 mice per phenotype) (B) UMAPs as in (A); red indicates normalized expression of selected genes. (C) UMAP as in (A) indicates normalized CD39 protein expression with yellow asterisks highlighting clusters with increased CD39 expression in γδ^HYP^ mice. (D) The frequency of CD39^hi^ cells among each cluster is shown. (E) Pseudobulk analysis was performed on CD39^neg^, CD39^int^, and CD39^hi^ cells (C4 and C5 combined) from both WT and γδ^HYP^ mice. Balloon plots representing the average expression and frequencies of selected surface receptors, immune signaling genes, transcriptional regulators, and effector molecules in γδ IELs from (A). (F) MFI of Aiolos among CD39-expressing γδ IEL populations in WT and γδ^HYP^ mice. n=5-6. (G) Balloon plot representing the average expression and frequencies of selected genes downstream of IL-15 signaling in CD39^int^ and CD39^hi^ γδ IELs from WT and γδ^HYP^ mice. Statistical analysis: ± 1.5-fold change, adjusted p-value of 0.05. (F) two-way ANOVA with Tukey’s post hoc test.

We next pooled clusters 4 and 5 from both phenotypes to interrogate the transcriptome of all γδ IELs based on CD39 protein expression. From these analyses, we were able to identify 37 DEGs between CD39^neg^ and CD39^int^ cells, 40 DEGs between CD39^int^ and CD39^hi^ cells, and 149 DEGs between CD39^neg^ and CD39^hi^ γδ IELs (Supplementary Table 1). Our analysis revealed a gradient of gene expression that correlated positively or negatively with CD39 expression (Fig. 2E). As indicated by unsupervised clustering, CD39^neg^ γδ IELs show elevated expression of *Tcf7*, which is consistent with reduced effector function and a stem-like or memory phenotype^33, 34^. Consistent with this, CD39^neg^ γδ IELs have higher expression of *Thy1*, *Id3*, *Ltb*, *Cd160*, and *Xcl1*, suggesting that CD39^neg^ γδ IELs are more naïve-like relative to those that are CD39^pos^. In contrast, CD39 expression positively correlated with the expression of inhibitory receptors such as *Lag3*, *Cd200r2*, *Cd200r4*, *Klra5*, *Cd8a* and *Fcer1g*.

Indicative of a more tissue-adapted phenotype, CD39^hi^ γδ IELs exhibit increased expression of genes downstream of TCR and IL-15 signaling, such as *Il2rb*, *Tyrobp*, *Tnfrsf9*, *Pik3cd*, *Pim1*, *Gzma* and *Gzmb*. Although we observed increased *Ahr* expression by bulk RNAseq (Fig. 1D), *Ahr* transcript was not markedly affected by CD39 expression (Fig. 2E). Many of the genes upregulated in CD39^hi^ γδ IELs overlap with those expressed by *Tcf7*^neg^ *Il2rb*^hi^ natural IELs, a population that expanded after genetic deletion of *Ikzf3* (Aiolos)^34^. IELs from Aiolos-deficient mice were hyperresponsive to IL-15 yet Aiolos expression is similar across CD39 populations and phenotypes (Fig. 2F), suggesting that Aiolos may restrain cytotoxicity in CD39^hi^ γδ IELs.

Next, we investigated whether the γδ^HYP^ phenotype influences the transcriptional program of CD39^pos^ γδ IELs by comparing CD39^int^ and CD39^hi^ γδ IELs from WT and γδ^HYP^ mice. 370 DEGs were identified between CD39^int^ WT and γδ^HYP^ IELs, whereas 110 DEGs were found between CD39^hi^ γδ IELs from each phenotype (Supplementary Table 2). Several genes associated with IL-15 signaling were upregulated based on the phenotype and magnitude of CD39 expression (Fig. 2G). However, *Klra5*, *Pim1* and *Bcl2* show a striking increase in both the mean expression and fraction of expressing cells within the given subset in γδ^HYP^ mice.

These findings highlight IL-15 signaling as a potential upstream regulator of CD39 that may be further amplified in γδ^HYP^ mice.

### Elevated IL-15 signaling leads to the upregulation of CD39 in γδ^HYP^ IELs

We next explored the relationship between IL-15 signaling and CD39 on γδ^HYP^ IELs and found that IL-15RC was increased in γδ^HYP^ jejunal lysates compared to WT (Fig. 3A). Further, CD122 (IL-2Rβ) expression correlated with increased CD39 in γδ IELs, independent of the γδ^HYP^ phenotype (Fig. 3B). While the amount of cell surface CD122 did not vary widely based on CD39 expression, CD122 was slightly reduced among γδ IELs that expressed no or intermediate amounts of CD39 in γδ^HYP^ mice relative to WT (Fig. 3C). To determine whether IL-15 is capable of inducing CD39 expression, sorted WT CD39^neg^ and CD39^pos^ γδ IELs were stimulated *in vitro* with IL-15RC. Whereas low IL-15RC had minimal effect, exposure to a higher concentration was sufficient to drive CD39 expression in CD39^neg^ γδ IELs and further enhance its expression in CD39^pos^ γδ IELs (Fig. 3D,E). Blocking IL-15 signaling *in vivo* reduced the frequency of CD39^hi^ γδ IELs only in γδ^HYP^ mice (Fig. 3F,G). These findings highlight the critical role of IL-15 signaling in promoting γδ IEL CD39 expression, and suggests that increased IL-15RC within the mucosa further amplifies CD122-mediated CD39 expression in γδ^HYP^ IELs.

**Figure 3:**
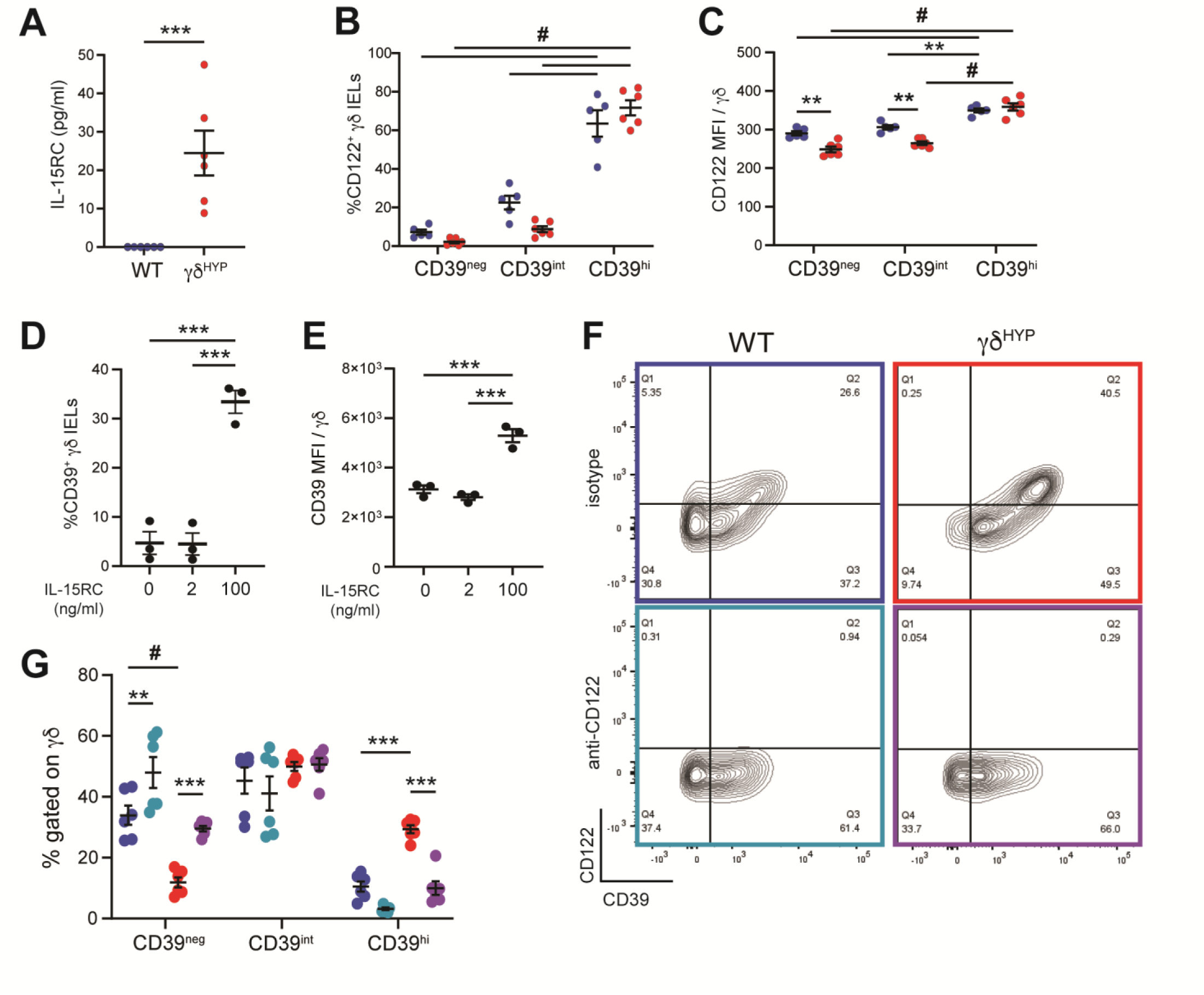
Enhanced IL-15 signaling leads to the upregulation of CD39 in γδ^HYP^ IELs. (A) ELISA of IL-15RC in WT and γδ^HYP^ jejunum. (B) Frequency and (C) MFI of CD122^+^ γδ IELs among CD39 populations. WT CD39^neg^ or CD39^pos^ γδ IELs were sorted and stimulated *in vitro* with 2 or 100 ng/ml IL-15RC for 72 h. (D) The frequency of CD39^neg^ γδ IELs that became CD39^pos^ after stimulation. (E) CD39 MFI of sorted CD39^pos^ γδ IELs following IL-15RC treatment. WT and γδ^HYP^ mice were injected i.p. daily with isotype control or anti-CD122 for 72 h. (F) Representative flow plots showing CD122 and CD39 expression on γδ IELs and (G) frequency of CD39^neg^, CD39^int^, and CD39^hi^ γδ IELs. All data shown as mean ± SEM from at least 2 independent experiments. (A-C,F,G) Each data point represents an individual mouse. n=5-6. (D,E) γδ IELs were pooled from multiple mice; each data point represents an individual replicate. n=3. Statistical analysis: (A) Unpaired student’s t-test, (B,C,G) two-way ANOVA with Tukey’s post hoc test, (D,E) one-way ANOVA with Tukey’s post hoc test. **P<0.01, ***P<0.001, #P<0.0001.

### TCR signaling in γδ^HYP^ IELs upregulates CD122 and CD39 expression

Our transcriptomic analyses indicate that TCR signaling may be activated in γδ^HYP^ IELs (Fig. 1D, 2E), therefore we asked if TCRγδ activation contributed to CD39 upregulation in phenotypic mice. Consistent with prior antigen exposure *in vivo*, surface TCRγδ was decreased on both CD39^neg^ and CD39^int^ γδ^HYP^ IELs relative to those from WT mice and CD39^hi^ γδ^HYP^ IELs (Fig. 4A,B). Notably, minimal TCR internalization was observed in WT mice or CD39^hi^ γδ^HYP^ IELs, suggesting that these cells are not responding a TCR signal. We next asked if TCR activation *in vitro* could influence CD122 and CD39 expression. Whereas IL-15RC was sufficient to induce CD39 expression and a modest increase in CD122 on CD39^neg^ γδ IELs, when combined with a TCR agonist, the expression of both proteins was substantially enhanced (Fig. 4C-E). Antibody-mediated blockade of TCR signaling *in vivo* resulted in TCR internalization in 30-40% of γδ IELs (Fig. 4F). Thus, we compared CD39 and CD122 expression between WT and γδ^HYP^ IELs that had internalized the TCR versus those that had not and found that abrogating the TCRγδ signal reduced the frequency of CD122 expression across CD39 populations in both WT and γδ^HYP^ IELs (Fig. 4G). Strikingly, impaired TCR signaling only decreased the frequency of CD39^hi^ γδ IELs in γδ^HYP^ mice (Fig. 4H). Taken together, our data demonstrate that increased TCR activation among CD39^neg^ γδ^HYP^ IELs promotes the expression of CD122 that subsequently drives CD39 upregulation.

**Figure 4.**
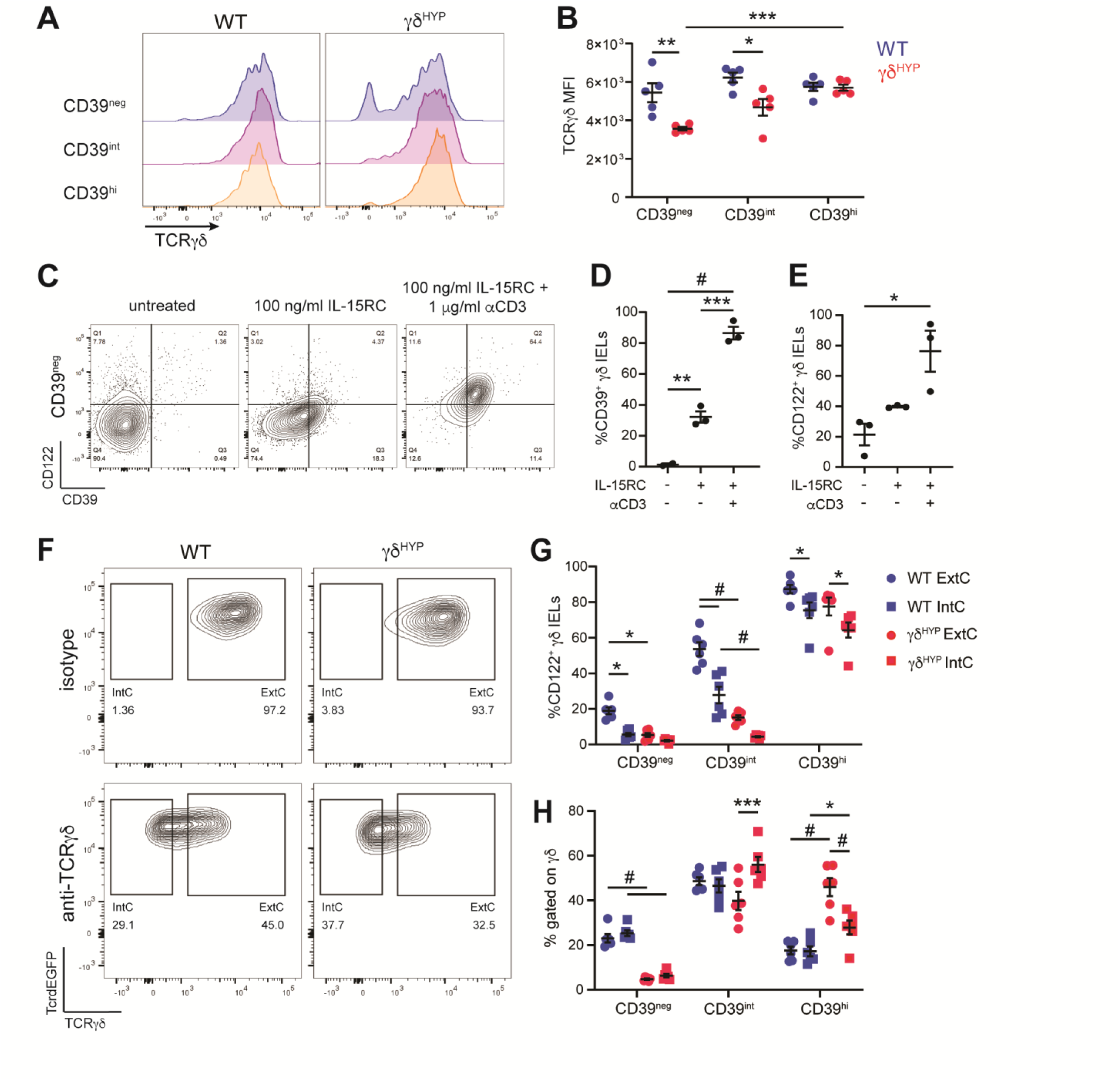
TCR signaling in γδ^HYP^ IELs leads to the upregulation of CD122 and CD39. (A) Representative flow plots and (B) MFI of TCRγδ showing TCR internalization among CD39 γδ IEL populations from WT and γδ^HYP^ mice. WT CD39^neg^ γδ IELs were sorted and stimulated *in vitro* with 100 ng/ml IL-15RC in the presence or absence of anti-CD3 for 72 h. (C) Representative flow plots and frequency of (D) CD39^neg^ γδ IELs that became CD39^pos^ and (E) CD122^+^ γδ IELs following stimulation. WT and γδ^HYP^ mice were injected i.p. daily with either isotype control or anti-TCRγδ antibody for 48 h. (F) Representative flow plots showing intracellular (IntC) versus extracellular (ExtC) TCRγδ after antibody treatment. (G) Frequency of CD122^+^ γδ IELs among CD39 populations and (H) frequency of CD39^neg^, CD39^int^, and CD39^hi^ γδ IELs in γδ IELs with IntC vs ExtC TCR. All data shown as mean ± SEM from at least 2 independent experiments. (B,G,H) Each data point represents an individual mouse. n=5-6. (C-E) γδ IELs were pooled from multiple mice, each data point represents an individual replicate. n=3. Statistical analysis: (B,G,H) two-way ANOVA with Tukey’s post hoc test, (D,E) one-way ANOVA with Tukey’s post hoc test. *P<0.05, **P<0.01, ***P<0.001, #P<0.0001.

### γδ^HYP^-associated microbiota sustains CD39^hi^ γδ IELs through increased CD122 expression

To determine if the increase of CD39 on γδ IELs requires the presence of γδ^HYP^-associated commensals, mice were administered broad-spectrum antibiotics (Abx) for 6 weeks to ensure complete turnover of the IEL compartment^35^. Abx treatment reduced the frequency of CD122^+^ CD39^+^ γδ IELs in both phenotypes, yet the relative proportion of CD39^hi^ γδ IELs was only decreased in γδ^HYP^ mice (Fig. 5A-C) mirroring our findings following TCRγδ blockade (Fig. 4G,H). Notably, neither the levels of IL-15RC in jejunal lysates nor the total number of γδ IELs were affected by depletion of the microbiota (Fig. 5D,E). From these data, we conclude the following: (1) continuous exposure to the γδ^HYP^-associated microbiota is required to activate the TCRγδ and promote the downstream expression of CD122 and CD39, and (2) sustained IL-15RC levels may be sufficient to support γδ^HYP^ IEL proliferation despite depletion of commensal bacteria.

A small subset of γδ IELs express a polyspecific TCR that can recognize Cy3 and structurally similar molecules, including indoles produced by Gram-positive bacteria^22^. To determine whether the activation of polyspecific TCRs is responsible for the expansion of CD39^hi^ γδ IELs in γδ^HYP^ mice, we treated mice with either vancomycin or neomycin to specifically deplete Gram-positive or-negative bacteria, respectively.

Administration of either antibiotic alone had no effect on CD39 expression on γδ^HYP^ IELs (Fig. 5F), indicating that indoles are unlikely the primary contributor to TCRγδ activation in phenotypic mice. Thus, the putative TCR ligand generated by γδ^HYP^-associated microbiota to drive γδ IEL CD122 and CD39 expression remains an area for future investigation.

**Figure 5.**
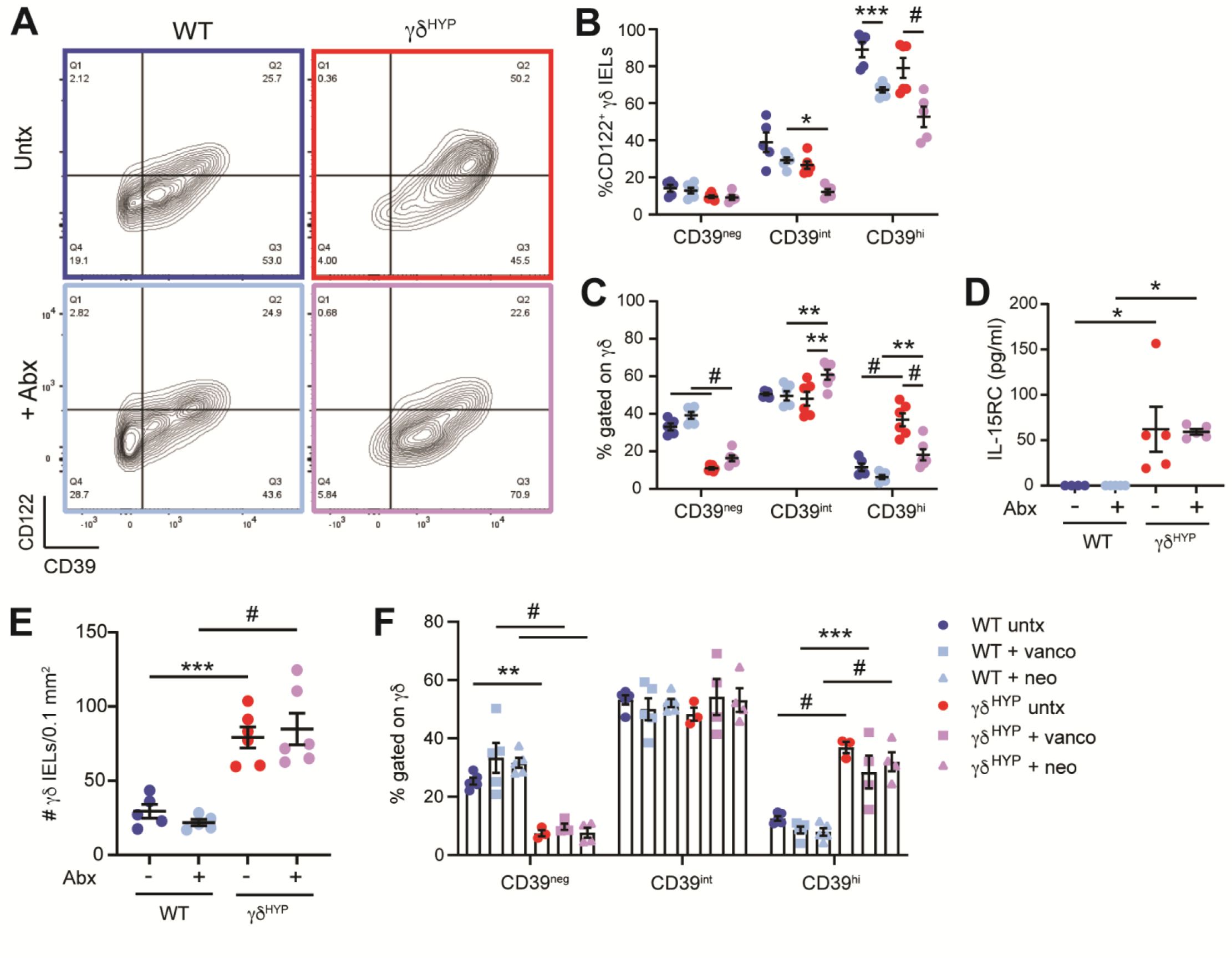
γδ^HYP^-associated microbiota sustains CD39^hi^ γδ IELs through increased CD122 expression. Mice were administered broad-spectrum antibiotics (Abx) in drinking water for 6 weeks. (A) Representative flow plots and frequency of (B) CD122^+^ γδ IELs among CD39 populations and (C) CD39^neg^, CD39^int^, and CD39^hi^ γδ IELs. (D) ELISA of IL-15RC in jejunal lysates. (E) Morphometric analysis of jejunal γδ IELs in untreated (untx) and Abx-treated mice. (F) WT and γδ^HYP^ mice were administered either vancomycin (vanco) or neomycin (neo) in drinking water for 6 weeks and the frequency of CD39 γδ IEL populations is shown. All data shown as mean ± SEM from at least 2 independent experiments. (B-F) Each data point represents an individual mouse. n=4-6. Statistical analysis: (B,C,F) Two-way ANOVA with Tukey’s post hoc test, (D,E) one-way ANOVA with Tukey’s post hoc test. *P<0.05, **P<0.01, ***P<0.001, #P<0.0001.

### Amplified IL-15 signaling promotes the survival of CD39^hi^ γδ^HYP^ IELs

We next investigated the extent to which the elevated mucosal IL-15RC may influence the maintenance of γδ IELs in phenotypic mice. Morphometric analysis revealed that treatment with anti-CD122 substantially reduced γδ IEL number in γδ^HYP^ mice to levels observed in WT (Fig. 6A). Since the frequency of CD39^hi^ γδ^HYP^ IELs was diminished in response to CD122 blockade *in vivo* (Fig. 3F,G), we hypothesized that IL-15 signaling may not only promote γδ IEL proliferation, but also prolong CD39^hi^ γδ^HYP^ IEL survival. In support of this, we observed increased Bcl-2 expression in CD39^hi^ γδ IELs that was further enhanced in γδ^HYP^ mice (Fig. 6B,C). Consistent with our *in vivo* findings, the addition of anti-CD122 *in vitro* resulted in the loss of CD39^hi^ γδ IELs (Fig. 6D), which may be a result of reduced Bcl-2 expression in CD39^pos^ γδ IELs among both phenotypes (Fig. 6E). Thus, increased mucosal IL-15RC enhances the long-term survival of CD39^hi^ γδ^HYP^ IELs by regulating Bcl-2 expression.

**Figure 6.**
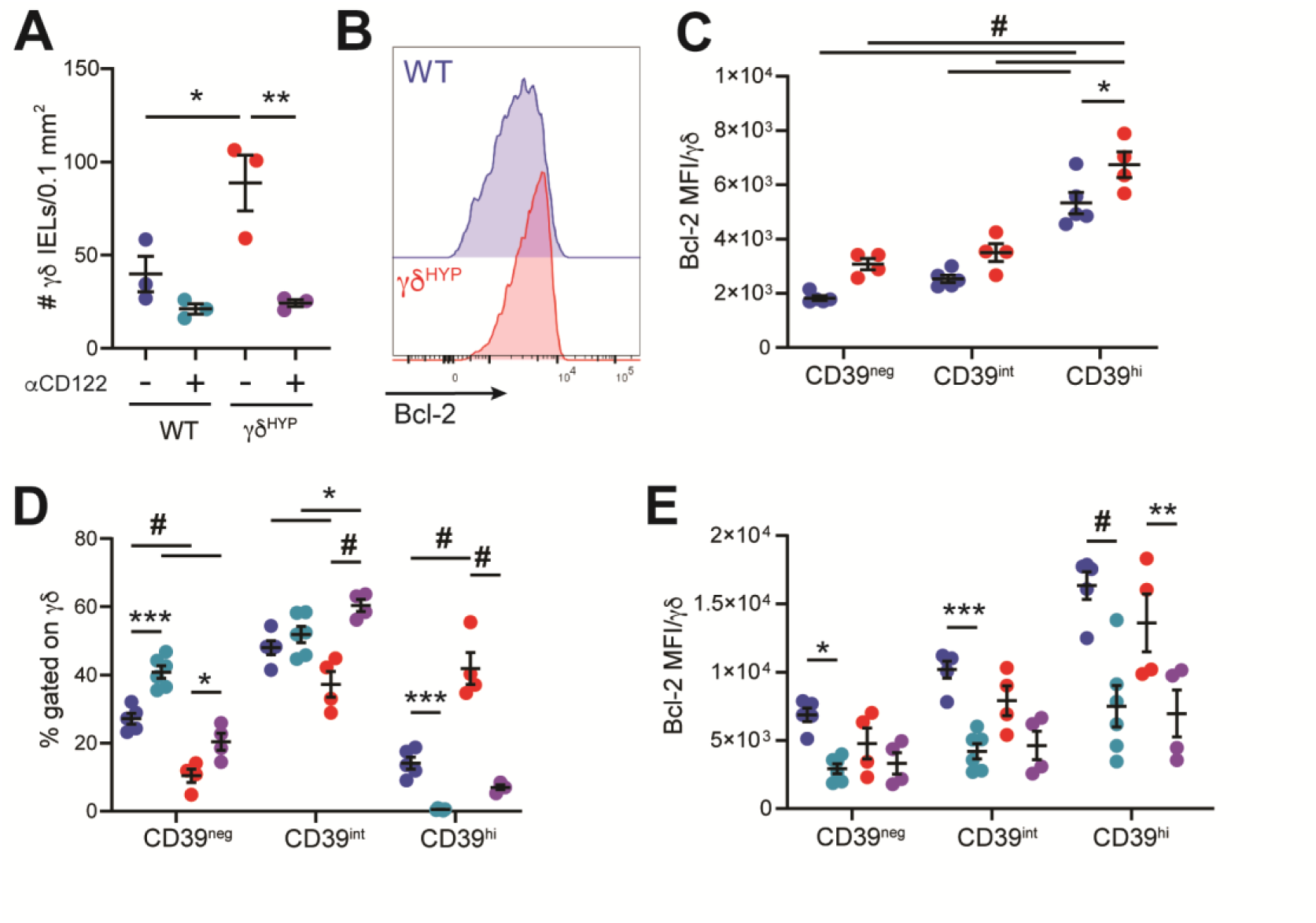
Amplified IL-15 signaling promotes the survival of CD39^hi^ γδ^HYP^ IELs. (A) WT and γδ^HYP^ mice were injected i.p. daily with isotype control or anti-CD122 for 72 h. Morphometric analysis of γδ IELs in WT and γδ^HYP^ jejunum. (B) Representative flow plot and (C) MFI of Bcl-2 among CD39 γδ IEL populations. WT and γδ^HYP^ IELs were isolated and cultured *in vitro* with 100 ng/ml IL-15RC in the presence or absence of anti-CD122 for 24 h. (D) Frequency of CD39 γδ IEL populations and (E) Bcl-2 MFI among CD39^neg^, CD39^int^, and CD39^hi^ γδ IELs. All data shown as mean ± SEM from at least 2 independent experiments. Each data point represents an individual mouse. n=4-6. Statistical analysis: (A) One-way ANOVA with Tukey’s post hoc test, (C-E)) two-way ANOVA with Tukey’s post hoc test. *P<0.05, **P<0.01, ***P<0.001, #P<0.0001.

### Elevated metabolism in γδ^HYP^ IELs supports protein synthesis

In addition to enhancing cell survival, IL-15 also increases IEL metabolic capacity and protein translation^5^. The observed upregulation of genes associated with AA and nutrient uptake transporters in γδ^HYP^ IELs (Fig. 1E) led us to ask whether IL-15 signaling may similarly regulate metabolism and protein translation in these cells.

Single Cell ENergetIc metabolism by profiling Translation inHibition (SCENITH^TM^)^36^ allows for determination of the cellular dependence for glucose and mitochondrial metabolism by assessing changes in protein synthesis in response to metabolic inhibitors. We found that the bioenergetic profile of γδ IELs at their metabolic peak^37^ was similar between the two phenotypes (Fig. 7A). Protein translation was higher in γδ^HYP^ IELs following activation compared to WT, which was abrogated in the presence of either metabolic inhibitor (Fig. 7B). Thus, while the bioenergetic profile of γδ^HYP^ IELs are similar to WT, these cells may retain increased metabolic capacity to support protein synthesis once stimulated.

**Figure 7.**
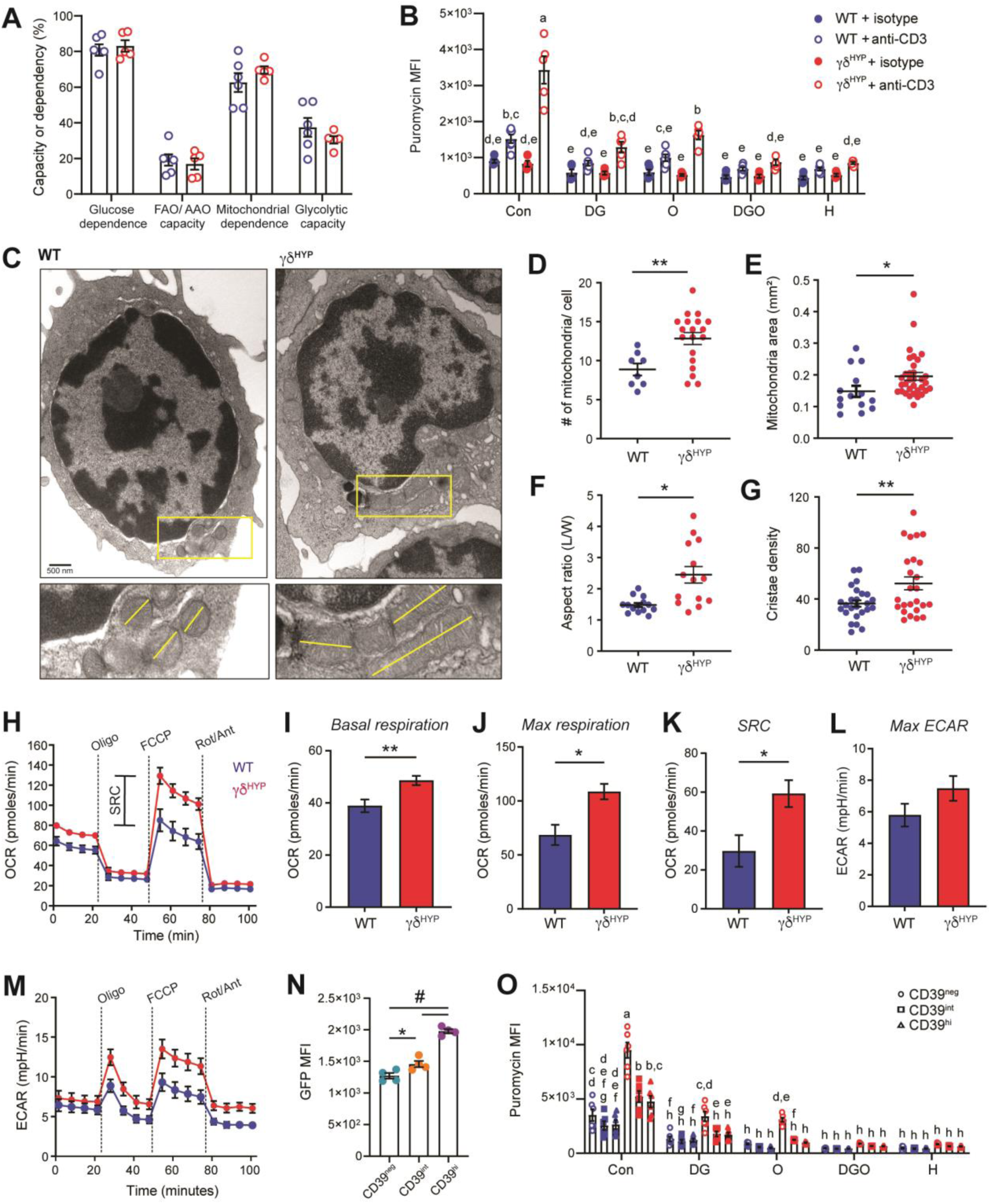
Elevated metabolism in γδ^HYP^ IELs supports protein production. SCENITH^TM^ was performed on γδ IELs isolated from WT and γδ^HYP^ mice injected i.p. with isotype control or anti-CD3 for 16 h. (A) Percent capacity or dependency of metabolic pathways upon stimulation and (B) puromycin MFI in response to DG, O, a combination of both (DGO), or harringtonine (H). (C) Transmission electron microscope images of sorted WT and γδ^HYP^ IELs. Scale bar= 500 nm. Yellow box shows inset and dashed yellow lines indicate the length of the mitochondrion. (D) Number of mitochondria per cell, (E) mitochondrial area (length x width, μm^2^), (F) aspect ratio (length/width) and (G) cristae density (number of cristae/area). Seahorse assays were performed on sorted, unstimulated WT and γδ^HYP^ IELs. (H) OCR, (I) basal respiration, (J) maximal respiration, and (K) SRC. (L) Maximal ECAR and (M) ECAR measured during OCR assay. (N) γδ IELs were isolated from Mito-QC mice and stained for CD39. GFP MFI of mitochondria among CD39 populations. (O) SCENITH^TM^ was performed on γδ IELs isolated from WT and γδ^HYP^ mice after 16 h anti-CD3 stimulation. Puromycin MFI among CD39^neg^, CD39^int^, and CD39^hi^ γδ IELs. All data shown as mean ± SEM from at least 2 independent experiments. (A,B,N,O) n=3-6. Each data point represents an individual mouse. (D-G) Cells were isolated and pooled from multiple mice. Each datapoint represents an average of mitochondria from individual cells. (H-M) Cells were pooled from multiple mice, each data point represents an individual replicate. Statistical analysis: (A,B,O) two-way ANOVA with Tukey’s post hoc test, different letters represent statistical significance. (D-G, I-L) Unpaired t-test, (N) one-way ANOVA with Tukey’s post hoc test. *P<0.05, **P<0.01, #P<0.0001.

IL-15 promotes mitochondrial fusion, cristae remodeling, and enhanced spare respiratory capacity (SRC) in conventional lymphocytes^38, 39^, thus we investigated whether γδ^HYP^ IELs are similarly affected. Transmission electron microscopy on sorted γδ IELs revealed an increase in mitochondrial number, area, aspect ratio, and cristae density in γδ^HYP^ IELs relative to WT (Fig. 7C-G), indicative of enhanced oxidative phosphorylation (OXPHOS). To determine whether the altered mitochondrial morphology in γδ^HYP^ IELs influenced cellular respiration, mitochondrial stress assays were performed to assess the oxygen consumption rate (OCR) with simultaneous measurement of the extracellular acidification rate (ECAR). While γδ^HYP^ IELs exhibit a small increase in basal OCR compared to WT, maximal respiration and SRC were substantially increased (Fig. 7H-K). Maximal ECAR was slightly elevated in both WT and γδ^HYP^ IELs following the addition of the mitochondrial uncoupler FCCP (Fig. 7L,M), reflecting the known link between glycolysis and OXPHOS in IELs^37^. This may explain why we fail to observe an overall change in the metabolic profile despite elevated OXPHOS in γδ^HYP^ IELs. From these data, we conclude that elevated mucosal IL-15RC may promote enhanced metabolic reserves in γδ^HYP^ IELs, yet basal respiration likely remains restricted to limit the potential for aberrant activation.

Prior transcriptomic analyses of human γδ IELs revealed an increase in genes associated with metabolic pathways in CD39^pos^ cells relative to those that were CD39^neg^ ^40^. Using mice that express GFP-labeled mitochondria^41^, we found that CD39 expression positively correlated with GFP intensity (Fig. 7N). Despite the increased mitochondrial mass in CD39^hi^ γδ IELs, metabolic capacity or dependency were similar among CD39-expressing γδ IEL populations (Supplementary Fig. 5A). We consistently observed enhanced protein synthesis across γδ^HYP^ IELs regardless of CD39 expression, yet the increase in translation was most prominent in CD39^neg^

γδ^HYP^ IELs (Fig. 7O). Notably, treatment with anti-CD3 *in vivo* did not have a substantial overall effect on CD39 frequency or expression in mice of either phenotype (Supplementary Fig. 5B,C). Since TCR activation induces CD122 to support proliferation in conventional T cells^42^, we next investigated the relationship between CD122, CD39 and cell proliferation. In response to anti-CD3, there is an increase in the frequency of CD122^pos^ CD39^neg^ γδ IELs that exhibit increased protein synthesis in both WT and γδ^HYP^ mice, but to a greater extent in the latter (Supplementary Fig. 5D-F). Of note, the translational capacity of CD122^pos^ CD39^hi^ γδ^HYP^ IELs was also increased compared to WT but to a reduced degree than in CD122^pos^ CD39^neg^ γδ IELs (Supplementary Fig. 5F). Taken together, we find that CD39^neg^ γδ^HYP^ IELs upregulate CD122 upon stimulation, and this increased responsiveness to IL-15 drives a higher basal bioenergetic profile to support cell growth and proliferation.

### CD39^hi^ γδ^HYP^ IELs display a reduced capacity for proliferation and pro-inflammatory cytokine production

We next asked whether CD39^hi^ γδ IELs exhibited differential proliferation or cytokine production in response to a TCR agonist *in vivo*. Consistent with the observed increase in protein synthesis (Supplementary Fig. 4F), proliferation was enhanced in CD39^neg^ γδ^HYP^ IELs whereas it was blunted in those with the highest CD39 expression (Fig. 8A). Interestingly, CD39^hi^ γδ IELs displayed reduced degranulation and produced negligible amounts of IFNγ or TNF in response to stimulation *in vitro* (Fig. 8B-D). Collectively, our findings suggest that the lack of intestinal pathology observed in γδ^HYP^ mice may be attributed the generation of a more mature, CD39^hi^ γδ IEL population with reduced cytotoxic potential.

**Figure 8:**
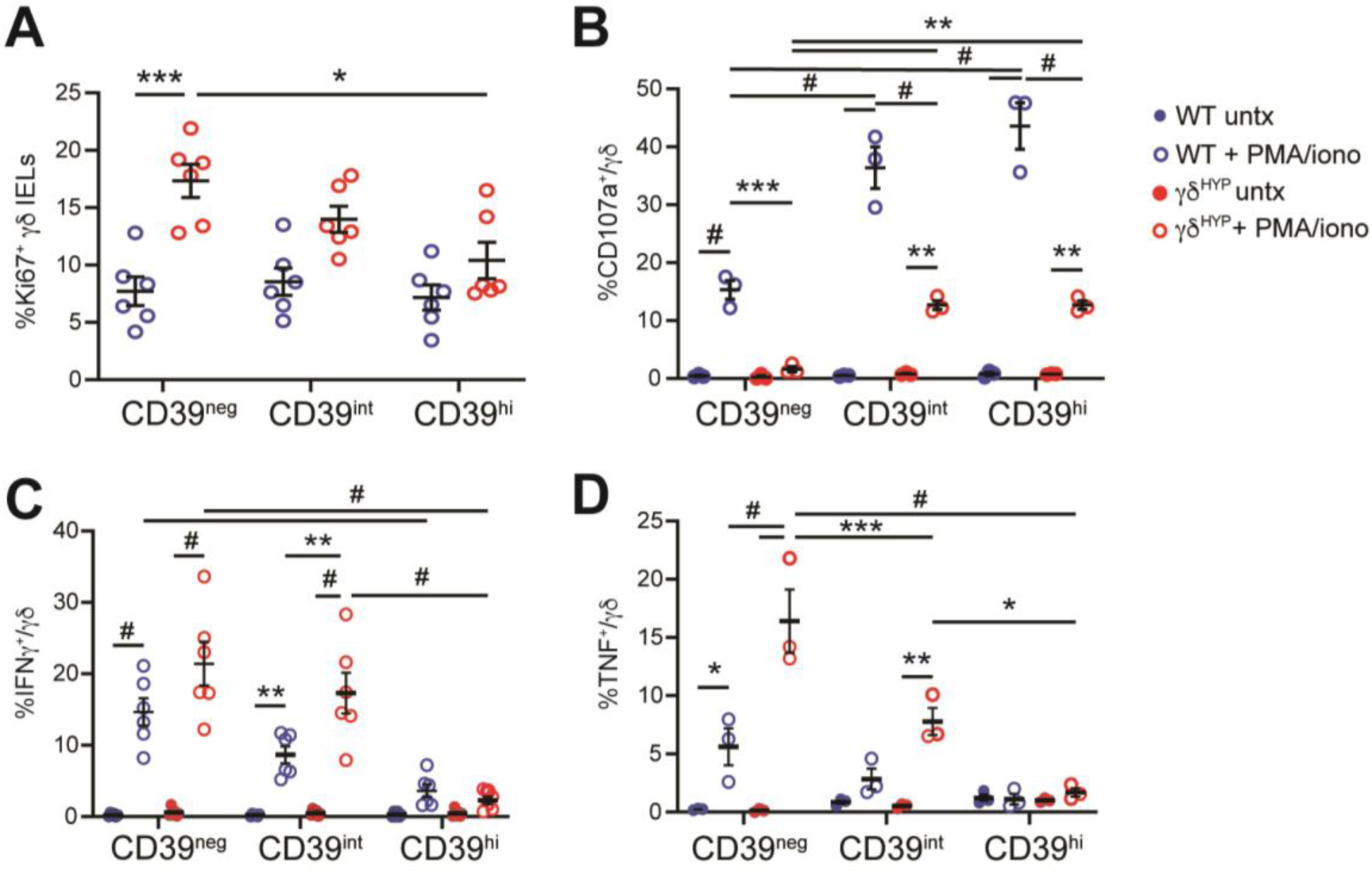
CD39^hi^ γδ^HYP^ IELs display reduced proliferative capacity and pro-inflammatory cytokine production. (A) WT and γδ^HYP^ mice were injected i.p with either isotype control or anti-CD3 for 16 h and Ki67^+^ γδ IELs among CD39^neg^, CD39^int^, and CD39^hi^ populations were assessed. Frequency of (B) CD107a^+^, (C) IFNγ^+^ and (D) TNF^+^ γδ IELs isolated from WT and γδ^HYP^ mice following *in vitro* 5 h stimulation with PMA/ionomycin. All data shown as mean ± SEM. Each data point represents an individual mouse. n=3-6. Statistical analysis: Two-way ANOVA with Tukey’s post hoc test. *P<0.05, **P<0.01, ***P<0.001, #P<0.0001.

## DISCUSSION

In this study, we identified a novel molecular mechanism by which an altered microbiota induces the expansion of a CD39^hi^ γδ IEL population with reduced cytotoxic capacity. Notably, the sustained exposure to γδ^HYP^-associated commensal bacteria is required for the maintenance of CD39^hi^ γδ^HYP^ IELs yet increases in IL-15RC expression or γδ IEL number do not return to WT levels in Abx-treated mice. These findings suggest that persistent TCR stimulation may be more critical for modulating CD39 expression and that the regulation of CD39 is uncoupled from cell proliferation. Our data demonstrate that CD39^neg^ CD122^+^ γδ IELs exhibit increased metabolic capacity and protein synthesis that likely supports enhanced proliferation among CD39^neg^γδ IELs. Although it is well known that PRR signaling promotes IL-15 expression by epithelial and immune cells^7–10^, the question remains: how are these high levels of mucosal IL-15 maintained once commensals are depleted? It is possible that exposure to the γδ^HYP^-associated microbiota induces long-term epigenetic modifications in IELs themselves or in IL-15-producing cells within the mucosa. Surprisingly little is known about the epigenetic regulation of γδ IELs^43^, and thus further investigation is required to determine whether transient exposure to the γδ^HYP^-associated microbiota can induce long-lasting changes to promote γδ IEL proliferation or maintenance within the epithelium.

In our prior characterization of this hyperproliferative phenotype, we found no evidence of clonal expansion among the γδ IEL population^23^, thus it is unlikely that a single antigen drives TCR activation to enhance CD39 expression. In support of this, we find that CD39 is upregulated across all three Vγ subsets in γδ^HYP^ mice. Since polyspecific γδ T cells recognize tyrosine residues^22^, we hypothesized that indoles produced by Gram-positive bacteria may stimulate the TCRγδ. However, no changes in CD39 expression among γδ IELs were observed following antibiotic-mediated depletion of Gram-positive bacteria. The ligands responsible for the activation of the murine TCRγδ largely remain unclear^44^, and therefore, further study is required to understand whether a γδ^HYP^-associated commensal ligand directly stimulates the TCR or if γδ IELs respond to a ligand displayed by epithelial cells or another mucosal immune cell.

Our data demonstrate that once immature or naïve-like CD39^neg^ γδ IELs are exposed to TCR activation and IL-15 in the mucosa, these cells upregulate CD122 and transition to a more mature, tissue-adapted CD39^hi^ population. The transcriptomic signature of CD39^hi^ γδ IELs indicates that these cells may be less responsive to TCR signals due to the upregulation of genes encoding inhibitory receptors such as *Lag3, Tyrobp* and *Fcer1g*, the last of which may ultimately attenuate the TCR signalosome^32^. This reduced activation state may explain (1) why TCR signaling is decreased in CD39^hi^ γδ IELs and (2) the substantial lack of cytokine production in these cells. Notably, TCR blockade leads to a decrease in CD122^+^ γδ IELs in both phenotypes yet we only observe a reduction in CD39 expression in γδ^HYP^ mice. Given that the TCR is exclusively activated in CD39^neg^ and CD39^int^ γδ^HYP^ IELs, it is possible that the absence of TCR signaling following receptor internalization prevents the maturation of γδ^HYP^ IELs and curtails CD39 upregulation. Alternatively, perturbation of IL-15 signaling through TCR downregulation may reduce the longevity of CD39^hi^ γδ^HYP^ IELs. Despite the low degree of TCRγδ responsiveness at steady-state^32, 45^, our data highlight a critical role for TCR stimulation in the expansion and maintenance of CD39^hi^ γδ^HYP^ IELs. IEL growth and proliferation are supported by increased nutrient uptake, SRC, and protein production; these processes are enhanced both in response to IL-15^5^ and in γδ^HYP^ IELs. Our observation of increased mitochondrial mass in CD39^hi^ γδ IELs fits with the known role of IL-15 in promoting T cell mitochondrial biogenesis^38^. Further, previous reports indicate that metabolic function correlates with CD39 expression in γδ IELs isolated from pediatric inflammatory bowel disease (IBD) patients^40^. It is worth noting that high levels of mucosal IL-15 are typically associated with increased licensing of IEL cytotoxicity; however, we observed the opposite in the γδ^HYP^ phenotype. In fact, we identified multiple layers of regulation that function to restrain γδ IEL activation in response to stimulus. For example, basal respiration remains low despite a clear increase in SRC in γδ^HYP^ IELs. Although there was a similar expansion of *Tcf7*^neg^ *Il2rb*^hi^ IELs in Aiolos-deficient mice^34^, CD39^hi^ γδ IELs maintain Aiolos expression to prevent an exaggerated response to IL-15. Moreover, we found that CD39^hi^ γδ IELs downregulate *Tcf7* expression and produce little effector cytokine, contrary to previous reports showing that *Tcf7*^-^ γδ IELs exhibit increased effector programming relative to *Tcf7*^+^ γδ IELs^33, 34^. These discrepancies highlight that in addition to cell-intrinsic regulation, the local microenvironment and microbial-derived signals likely also contribute to restraining γδ IEL activation.

We have recently shown that CD39^+^ γδ IELs exhibit immunosuppressive capacity^30^, demonstrating that CD39^+^ γδ IELs play a regulatory role in the gut. While further study is required to understand the role of CD39^+^ γδ IELs in maintaining mucosal homeostasis, the expansion of CD39^hi^ IELs may explain the lack of pathology in γδ^HYP^ mice. Future studies will investigate the immunosuppressive capacity of CD39^+^ γδ IELs *in vivo* and determine if adenosine production by γδ IELs signals in an autocrine fashion to regulate γδ IEL activation, or by suppressing other immune cells in the IEL compartment. CD39 is also a marker of T cell exhaustion^46, 47^; however, the unique activation state of IELs makes it difficult to determine whether these cells are truly exhausted despite a common proteomic signature^32^. Increased TCR stimulation may contribute to the upregulation of exhaustion markers in γδ^HYP^ IELs, or this could simply reflect γδ IEL maturation within the epithelium. Conversely, γδ^HYP^ IELs exhibit increased proliferation and elevated metabolic capacity in response to stimulation, which is inconsistent with an exhausted phenotype. While exhaustion may indicate a loss of pro-inflammatory effector function, this change in activation state can be viewed as a functional switch toward a more regulatory phenotype. This is particularly beneficial in the context of autoimmunity^27^, and thus we suggest that in combination with other local cues, TCRγδ activation may promote the maturation and enhance the suppressive capacity of γδ IELs.

Crohn’s disease (CD) and ulcerative colitis (UC) are relapsing and remitting inflammatory diseases of the gastrointestinal tract. Patients with IBD exhibit a reduction in γδ IEL number^48, 49^, yet this loss is reversible as γδ IELs are present in healed ulcerations^50^. We recently showed that this decrease in γδ IELs precedes the onset of chronic CD-like ileitis, with the frequency of CD39^+^ γδ IELs reduced a month prior to the initiation of ileal inflammation^30^. These data suggest that CD39^+^ γδ IELs are critical for mucosal homeostasis and that loss of this population may lead to a loss of immunological tolerance and disease development. A previous study showed that treatment with dipyridamole, a modulator of cyclic AMP signaling, increased CD39 expression in CD8 T cells and alleviated both experimental and pediatric colitis^40^. These findings suggest that identifying the microbes or ligands involved in amplifying CD39 expression, in addition to boosting γδ IEL number, may provide a novel therapeutic strategy to maintain mucosal homeostasis and prevent disease relapse in patients with IBD.

## MATERIALS AND METHODS

### Animals

Mice maintained on a C57BL/6 background, housed under specific pathogen free conditions, and provided autoclaved commercial rodent 5010 diet (LabDiet) and tap water. All mice were housed in a room with standard 12 h light-dark cycle and humidity and temperature were monitored. TcrdH2BEGFP (TcrdEGFP) mice were provided by Bernard Malissen (INSERM)^51^ and Mito-QC mice were obtained from Prof. Ian Ganley (University of Dundee). γδ^HYP^ mice were generated as F2 WT mice following vertical transfer of microbiota as previously reported^23^. Mice of both sexes were used between 8 to 12 weeks of age, except for *in vitro* experiments in which 15-20-week-old mice were used to obtain sufficient numbers of CD39^neg^ γδ IELs. Mice were injected with 0.45 mg anti-CD122 (TM-β1, BioXCell) or IgG2b isotype control (LTF-2, BioXCell) i.p daily for 3 days prior to the experiment. Alternatively, mice were treated with 200 μg anti-TCRγδ (UC7-13D5) blocking antibody or Armenian Hamster IgG (BioXCell, BE0091) i.p. daily for 2 days prior to euthanasia. For SCENITH^TM^ experiments, mice were injected with either 25 μg anti-CD3 (145-2C11, BioXCell) or Armenian Hamster IgG for 16 h. Mice received broad-spectrum antibiotics in drinking water containing 400 μg/ml vancomycin (Hospira) and 200 μg/ml meropenem (Bluepoint Laboratories) for 6 weeks. For antibiotic treatments targeting Gram-positive and Gram-negative bacteria, mice received either 500 μg/ml vancomycin (Hospira) or 1 mg/ml neomycin (Sigma-Aldrich). All studies were conducted in an Association of the Assessment and Accreditation of Laboratory Animal Care-accredited facility according to protocols approved by Rutgers New Jersey Medical School Comparative Medicine Resources and the Icahn School of Medicine Center for Comparative Medicine and Surgery.

### Immunostaining

For immunostaining, 7 μm thick frozen jejunal sections were blocked in 10% normal goat serum and stained with rabbit anti-laminin antibody (Sigma-Aldrich), AlexaFluor 594 goat anti-rabbit (Thermo Fisher Scientific), phalloidin AlexaFluor 647 and Hoechst 33342 before mounting with Prolong Glass (Life Technologies). Fluorescence micrographs were captured on an inverted Dragonfly 620 microscope (Andor) equipped with a dual microlens spinning disk, Sona 4.6 back-illuminated sCMOS camera (Andor), PlanApo 20x/0.8 dry objective (Nikon). For quantification of γδ IELs, images were taken on an upright widefield Axioimager.Z2(M) microscope (ZEISS) equipped with a Axiocam 503 color camera (ZEISS) and PlanApo 20x/0.8 dry objective (ZEISS). The number of γδ IELs per 0.1 mm^2^ of jejunum was quantified using FIJI (NIH).

### Flow cytometry

IELs were isolated as previously described^23^. Briefly, mice were euthanized, and the small intestine was opened longitudinally after the removal of Peyer’s patches. One cm pieces were incubated in HBSS with 3 mM EDTA and 7.5% FBS (Thermo Fisher Scientific) for 1 h in a shaker at 37°C. Supernatants were collected and IELs were isolated using a 20/45/70% Percoll density gradient (Cytiva). IELs were stained with fixable viability dye eFluor 780 (eBioscience), anti-CD3 (145-2C11), anti-TCRβ (H57-597), anti-CD4 (RM4-5), anti-CD8α (53-6.7), anti-CD8β (YTS1567.7), anti-CD39 (Duha69), anti-CD73 (TY/11.8), anti-TCRδ (GL3) purchased from Biolegend, anti-Vγ1 (2.11, BD Biosciences), anti-Vγ4 (UC3-10A6) and anti-CD122 (TM-β1) from BD Biosciences, biotinylated anti-Vγ7 (GL7, provided by Rebecca O’Brien (National Jewish Health, Denver, CO)) followed by streptavidin-BUV737 (BD Biosciences). Intracellular staining was performed following fixation with Cytofix/Cytoperm (Becton Dickinson) prior to staining with anti-IFNγ (XMG1.2, BioLegend), anti-TNF (MP6-XT22, BD Biosciences), and anti-CD107a (1D4B, BioLegend). Alternatively, the eBioscience Foxp3 transcription factor kit was used to stain for anti-Bcl2 (BCL/10C4, BioLegend), anti-Ki67 (16A8, BioLegend), anti-Aiolos (8B2, BioLegend) or anti-puromycin AF647 (R4743L-E8, 1:50). Flow cytometry was performed on a LSR Fortessa or FACSymphony A5 SE Cell Analyzer (BD Biosciences) and the data was analyzed using FlowJo (v. 10.10.0; Tree Star). IELs were sorted to 95-98% purity based on GFP expression using a BD FACSAria Fusion sorter.

### Bulk RNA sequencing

γδ IELs were isolated from WT and γδ^HYP^ mice, sorted based on Vγ1 and Vγ7 TCR expression, and RNA was extracted and purified using a RNeasy mini kit (Qiagen). RNA libraries were prepared using NEBNext Ultra II RNA Library Prep Kit for Illumina by PolyA selection. cDNA was synthesized, purified, and subjected to end-prep reaction, adaptor ligation and barcoded using unique combination of forward (i7) and reverse (i5) index primers. The cDNA libraries were sequenced using NextSeq 500 instrument (Illumina, San Diego, CA). Subsequent fastq files were processed using Kallisto (version 0.46.1) with 100 bootstraps for raw read counts and an average fragment length of 100 bp (20 bp standard deviation), then indexed to the mouse (mm9) genome. Transcript abundance HDF5 binary files were imported with tximport and then processed using DESeq2 (version 1.32) for calculating FPKM normalization and for differential expression analysis in R. Genes for analysis were filtered for an average FPKM of >1. For analyses in which transcripts from Vγ1 and Vγ7 IELs were combined and assessed based on phenotype, raw data files were converted to 50 fastq files, demultiplexed using bcl2fastq 2.20 software, and analyzed using Partek Flow pipeline (Partek, PGS7.21.1119). In Partek, reads were aligned to the most recently annotated Mus musculus (mm39) genome using STAR aligner (version 2.7.8a) and assigned to genomic features using the Partek E/M annotation model and annotated with Ensembl transcript (release 104). Genes with a maximum read count ≤100 reads were filtered out and remaining reads were normalized to the median ratio. DESeq2 was used to determine differential expression between WT and γδ^HYP^ samples. Differential gene expression (DGE) was filtered based on a false discovery rate (FDR) < 0.05 and a fold change of ± 1.5. Gene set enrichment (GSE) used the 2023-09-13 Gene Ontology database (GO Consortium) to identify GO terms associated with up or downregulated DEGs with a p<0.05. Genes sets were filtered to contain less than 1,000 genes.

### CITE-seq

CD3^+^ IELs were FACS-sorted from WT and γδ^HYP^ mice and debris-free suspensions of >80% viability were each labeled with a distinct Totalseq C mouse hashtag antibody (HTO-1 to HTO-6, Biolegend) to assign a unique sample identifier. All samples were subsequently pooled in equal proportions and stained with anti-CD39 antibody conjugated with oligonucleotide tags (Biolegend), following the manufacturer’s instructions. Cells were encapsulated into Gel-Bead in Emulsions (GEMs) using the Chromium X platform (10x Genomics) with the 5’ gene expression (5’ GEX) V2 kit, with a targeted recovery of 20,000 cells per lane. Cells were lysed within the GEMs, and both RNA and protein information were captured during reverse transcription (RT) and cDNA amplification steps, following the manufacturer’s established protocols. The resulting libraries contain information on gene expression (GEX), hashtag (HTO) and cell surface protein (ADT) levels for each cell. Libraries were sequenced in paired-end mode on a NovaSeq instrument (Illumina) targeting a depth of 25,000 reads per cell for GEX library and 5000 reads per cell for HTO and ADT libraries. Raw fastq files were aligned to the mm10 reference genome (2020-A) using Cell Ranger multi v7.0.1 (10x Genomics).

Analysis of scRNA seq data was conducted in Python using Scanpy library^52^ with default parameters unless otherwise specified. Quality control and filtering steps were performed to remove low-quality cells and genes prior to analysis. Pre-processing including normalization, principal component analysis, and Leiden clustering were performed followed by UMAP for visualization. Leiden clusters were manually annotated as specific cell types based on expression of marker genes (*Cd4*, *Cd8a*, *Cd8b*, *Trdc*, *Ki67*). Cells were categorized into CD39^neg^, CD39^int^, and CD39^hi^ groups based on normalized CD39 expression thresholds (0.15 <= x <= 3).

γδ IELs (CD8αα TCRγδ *Tcf7*^pos^ and CD8αα TCRγδ *Tcf7*^neg^) were selected for further downstream analyses. DGE analysis was performed by aggregating raw counts into pseudobulk per sample, filtering out genes with fewer than 10 counts. The DESeq2 package^53^ was used for DGE analysis with WT as reference. Variance stabilizing transform (vst) was applied to the pseudobulk data using DESeq2 vst function. The top 500 genes were selected using median absolute dispersion score^54^. Finally, the data was z-normalized before creating clustermap using Seaborn library^55^.

### In vitro stimulations

IELs were isolated and plated at 4×10^5^ cells/ well in media with or without 40 ng/ml PMA and 4 μg/ml ionomycin (Sigma-Aldrich) for 5 h (37°C, 5% CO_2_). GolgiPlug (BD Biosciences) was added for the final 4 h prior to analysis of IFNγ, TNF, and CD107a. For *in vitro* CD122 blocking experiments, 1.5×10^5^ IELs were plated and cultured overnight in 100 ng/ml IL-15RC, after which 40 μg/ml anti-CD122 (TM-β1, BioLegend) was added for 24 h. For experiments investigating the role of IL-15 or TCR stimulation on CD39 expression, IELs were isolated, sorted based on CD39 expression, and plated at 1×10^5^ cells/well in the presence of 2 or 100 ng/ml IL-15RC (eBioscience) for 72 h. Alternatively, sorted WT CD39^neg^ γδ IELs were cultured with either 100 ng/ml IL-15RC in the presence or absence of 1 μg/ml anti-CD3 (2C11, BioLegend) for 72 h. Freshly isolated and fixed CD39^neg^ or CD39^pos^ γδ IELs served as untreated controls.

### ELISA

One cm pieces of jejunum were harvested and homogenized with a bead beater (Precellys 24 Touch Homogenizer, Bertin Technologies) in Bio-Plex Cell Lysis Buffer (Bio-Rad). Protein concentration was determined using the DC Protein Assay (Bio-Rad). Lysates were diluted to 1 mg and analyzed with an IL15/IL15R Complex Mouse Uncoated ELISA Kit (ThermoFisher Scientific) per manufacturer’s instructions.

*SCENITH*^TM^. The SCENITH^TM^ Kit containing all reagents and protocols were provided by R. Argüello (GammaOmics)^36^. IELs isolated from anti-CD3-or isotype-treated mice were plated at 4×10^5^ cell/well and incubated for 45 min at 37°C, 5% CO^2^ with control, 2-deoxyglucose (100 mM, Sigma-Aldrich), oligomycin (1 mM, Sigma-Aldrich) or both inhibitors. IELs were treated with harringtonine (2 mg/ml, Abcam) to inhibit protein translation as a positive control. 2x puromycin was simultaneously added at the beginning of treatment. After 15 minutes, 50 mM verapamil (Sigma-Aldrich) was added to the media to prevent the efflux of puromycin through ABC transporters. Puromycin incorporation was determined by flow cytometry, as described above. The capacity and dependency of metabolic pathways were calculated using formulas previously described^36^.

### Transmission Electron Microscopy

5×10^6^ sorted small intestinal GFP^+^ γδ IELs from WT and γδ^HYP^ mice were pelleted and fixed in 2.5% glutaraldehyde, 4% paraformaldehyde in 0.1 M cacodylate buffer and post-fixed in buffered 1% osmium tetroxide. Samples were subsequently dehydrated in a graded series of acetone and embedded in Embed812 resin. 90 nm thin sections were cut on a Leica UC6 ultramicrotome and stained with saturated solution of uranyl acetate and lead citrate. Images were captured with an AMT (Advanced Microscopy Techniques) XR111 digital camera at 80Kv on a Philips CM12 transmission electron microscope. The number of mitochondria, aspect ratio (length/width), mitochondrial area (μm^2^), and cristae density were quantified using Image J. Samples were processed and images were acquired at Rutgers Robert Wood Johnson Medical School Electron Microscopy Core Imaging Lab.

### Seahorse assay

Agilent Seahorse XFp Cell Mito Stress Test was run per the manufacturer’s instructions. Sorted γδ IELs isolated from WT and γδ^HYP^ mice were washed twice with XF medium (DMEM medium, pH 7.4 containing 25 mM glucose, 2 mM L-glutamine, 1 mM sodium pyruvate) and seeded at 3×10^5^ cells/well. Oxygen consumption rate (OCR) and extracellular acidification rate (ECAR) were measured in Seahorse XF HS Mini Analyzer (Agilent) under basal conditions as well as in response to 2 μM oligomycin, 2 μM FCCP, and 1 μM rotenone with 1 μM antimycin A (Agilent).

### Statistical analyses

Data are represented as the mean ± SEM and statistical analyses were performed using Prism (GraphPad, v10). Unpaired student’s t tests were used to compare between two conditions, whereas one-way or two-way ANOVA with Tukey’s post hoc test were used to compare between three or more conditions. For the transcriptomics studies, a single experimental replicate with 3-4 mice per group was performed. All other experiments were performed represent at least two independent experiments.

## Acknowledgments

Cell sorting was performed at the Rutgers NJMS Flow Cytometry and Immunology Core Laboratory supported by National Institute for Research Resources Grant (S10 RR027022) and the Mount Sinai Dean’s Flow Cytometry CoRE. We would like to thank Rajesh Patel at the Rutgers Core Imaging Lab for his assistance in preparing samples for electron microscopy. We also extend our gratitude to Zhihong Chen from the Mount Sinai Human Immune Monitoring Core for his assistance with the experimental design for CITE-seq experiments.

## Author contributions

S.A. designed and performed experiments and wrote the manuscript. L.J., A.F., A.L., N.S. and M.S. performed experiments and analyzed the data. H.K. and M.N. analyzed data. R.J.A. provided resources and advised the research. M.P.V, M.S., and K.S.L. provided resources and supervised the research. K.L.E. conceived the study, supervised the research and wrote the manuscript. All authors approved the final manuscript.

## Data and materials availability

All bulk RNA-seq data generated in this study have been deposited in GEO under accession number GSE292555. Single cell RNA-seq data are in the process of being deposited at the time of submission.

---

**Supplementary Figure 1.**
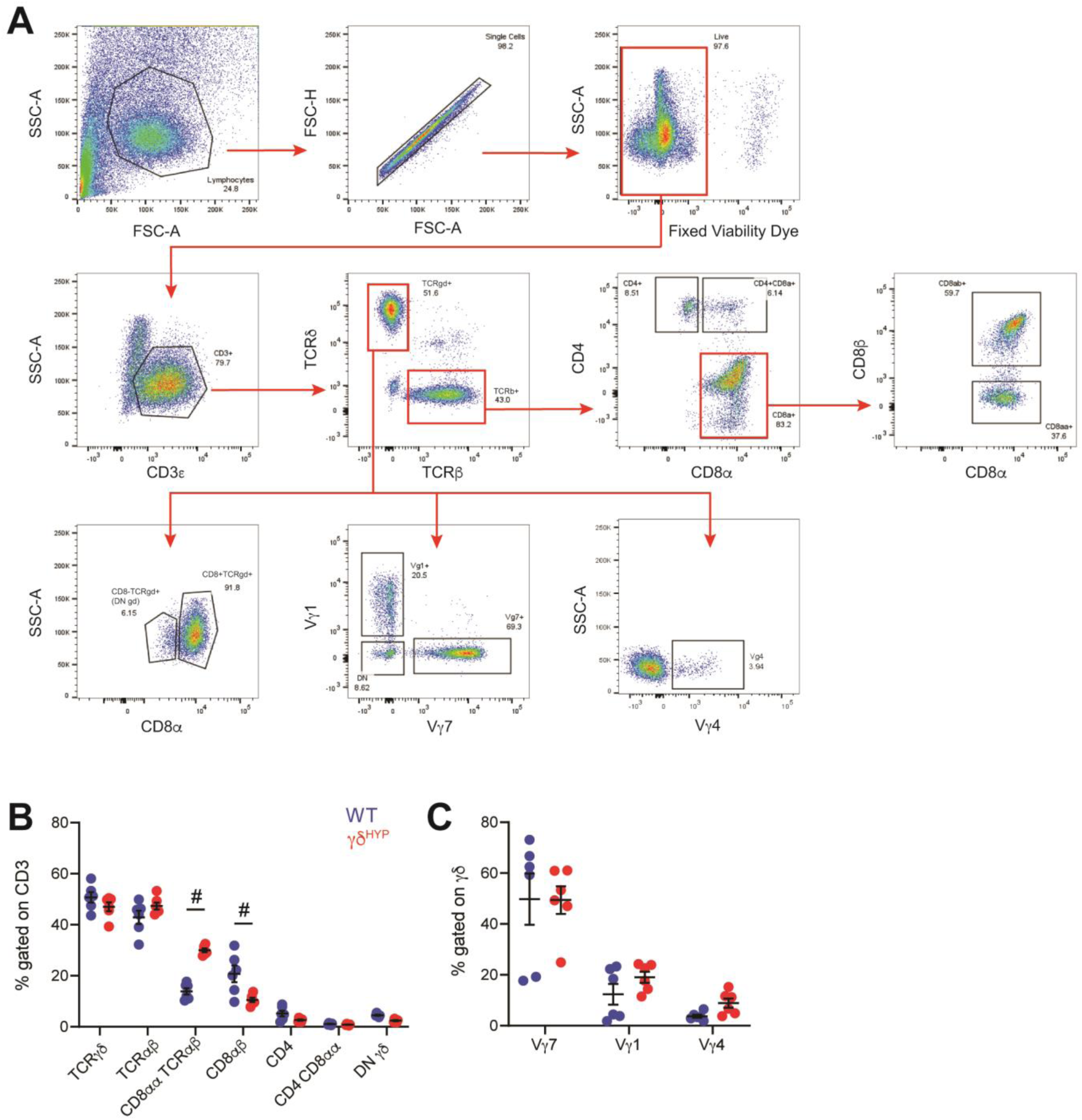
Natural IEL populations are expanded in γδ^HYP^ mice. (A) Gating strategy for IEL subsets. Frequency of (B) IEL populations gated on CD3+ and (C) Vγ subsets gated on TCRγδ^+^ IEls in WT and γδ^HYP^ mice. All data shown as mean ± SEM from at least 2 independent experiments. Each data point represents an individual mouse. n=6. Statistical analysis: two-way ANOVA with Tukey’s post hoc test #P<0.0001.

**Supplementary Figure 2:**
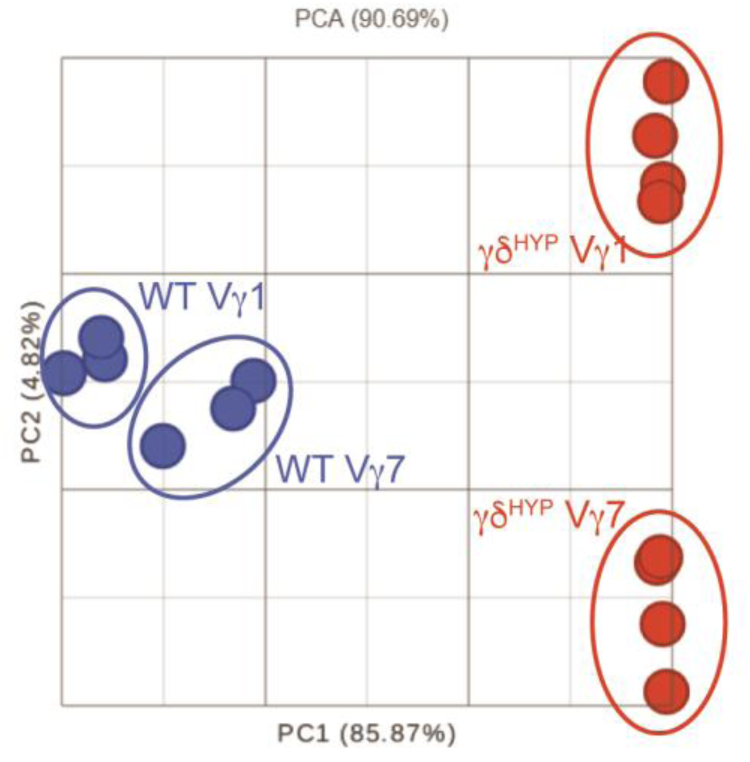
γδ IELs segregate based on the γδ^HYP^ phenotype. Principal component analysis of bulk RNA sequencing data of sorted Vγ1 and Vγ7 IELs isolated from WT or γδ^HYP^ mice. n=3-4.

**Supplementary Figure 3:**
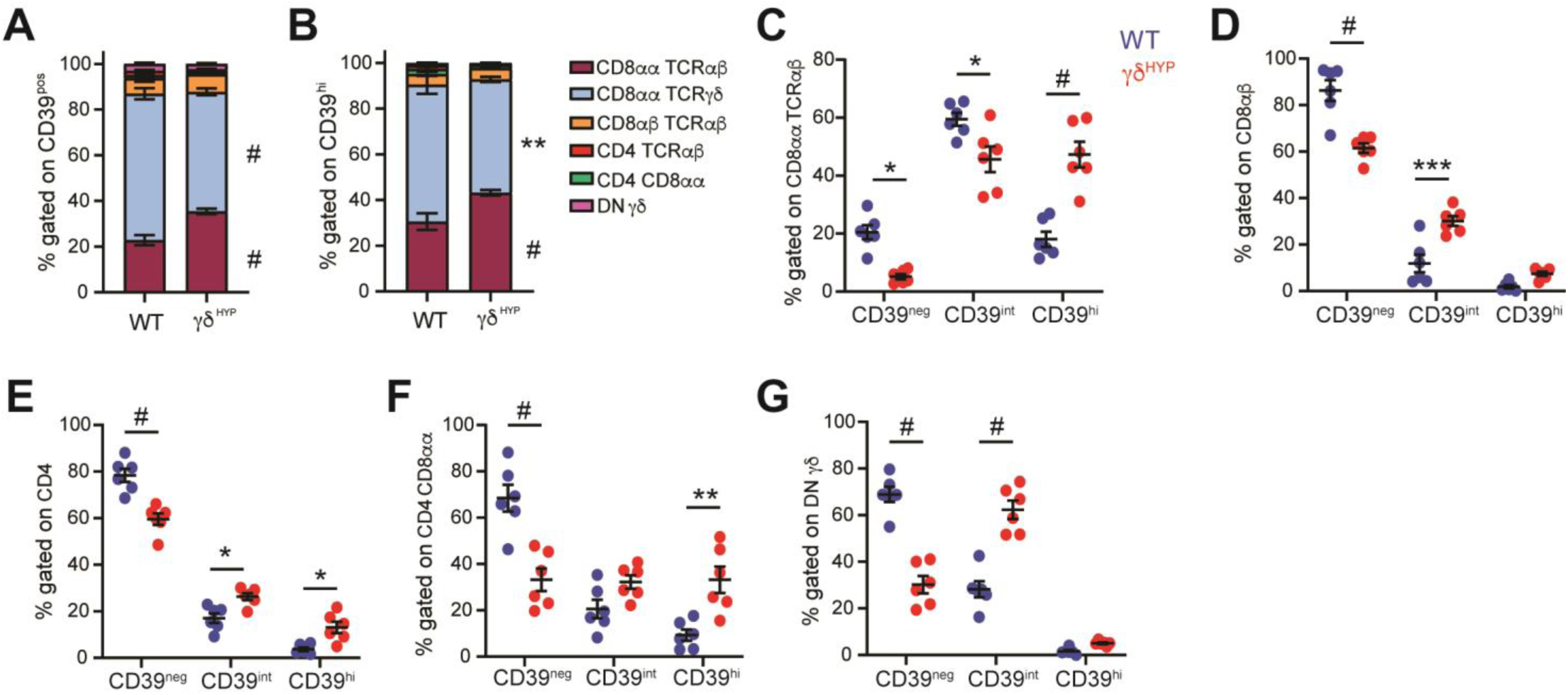
Natural IELs are the major contributors to CD39 expression in the IEL compartment. Frequency of IEL populations gated on (A) CD39^pos^ cells and (B) CD39^hi^ cells. Frequency of CD39^ne^^8^, CD39^iflt^, and CD39^hi^ cells gated on (C) CD8αα TCRαβ, (D) CD8αβ, (E) CD4, (F) CD4 CD8αα, and (G) DN γδ (CD8α TCRγδ) IELs. All data shown as mean ± SEM from at least 2 independent experiments. n=6. Each data point represents an individual mouse. Statistical analysis: two-way ANOVA with Tukey’s post hoc test. *P<0.05, **P<0.01, ***P<0.001, #P<0.0001.

**Supplementary Figure 4:**
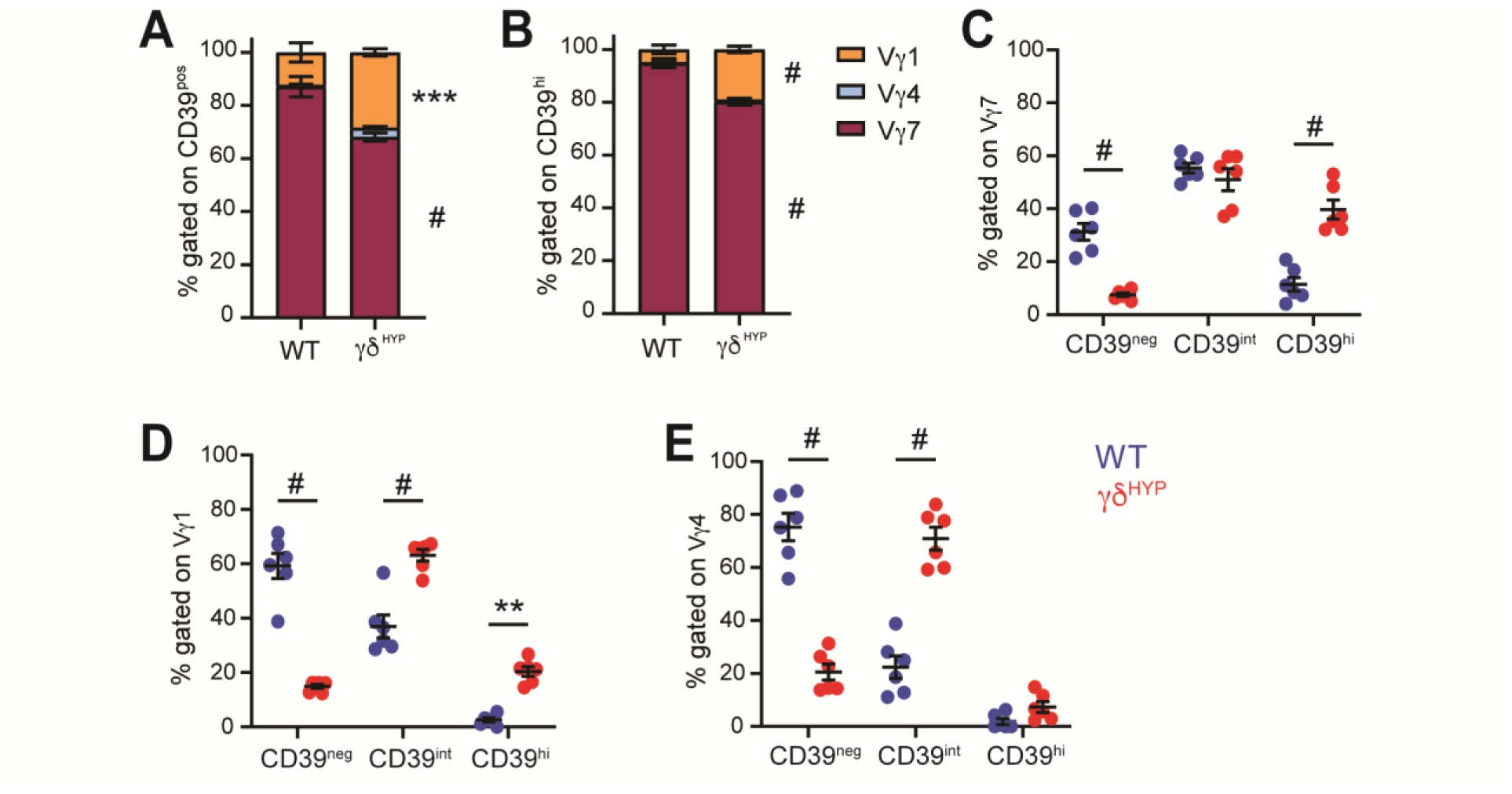
The majority of CD39^POS^ 78 IELs express Vy7 TCR. Frequency of Vy subsets gated on (A) CD39^p0S^ cells and (B) CD39^hl^ cells. Frequency of CD39^ne^^9^, CD39’^rt^, and CD39^hi^ cells gated on (C) Vγ7, (D) Vγ1 and (E) Vγ4 γ3 IELs. All data shown as mean ± SEM from at least 2 independent experiments. Each data point represents an individual mouse. n=6. Statistical analysis: two-way ANOVA with Tukey’s post hoc test. #P<0.0001.

**Supplementary Figure 5.**
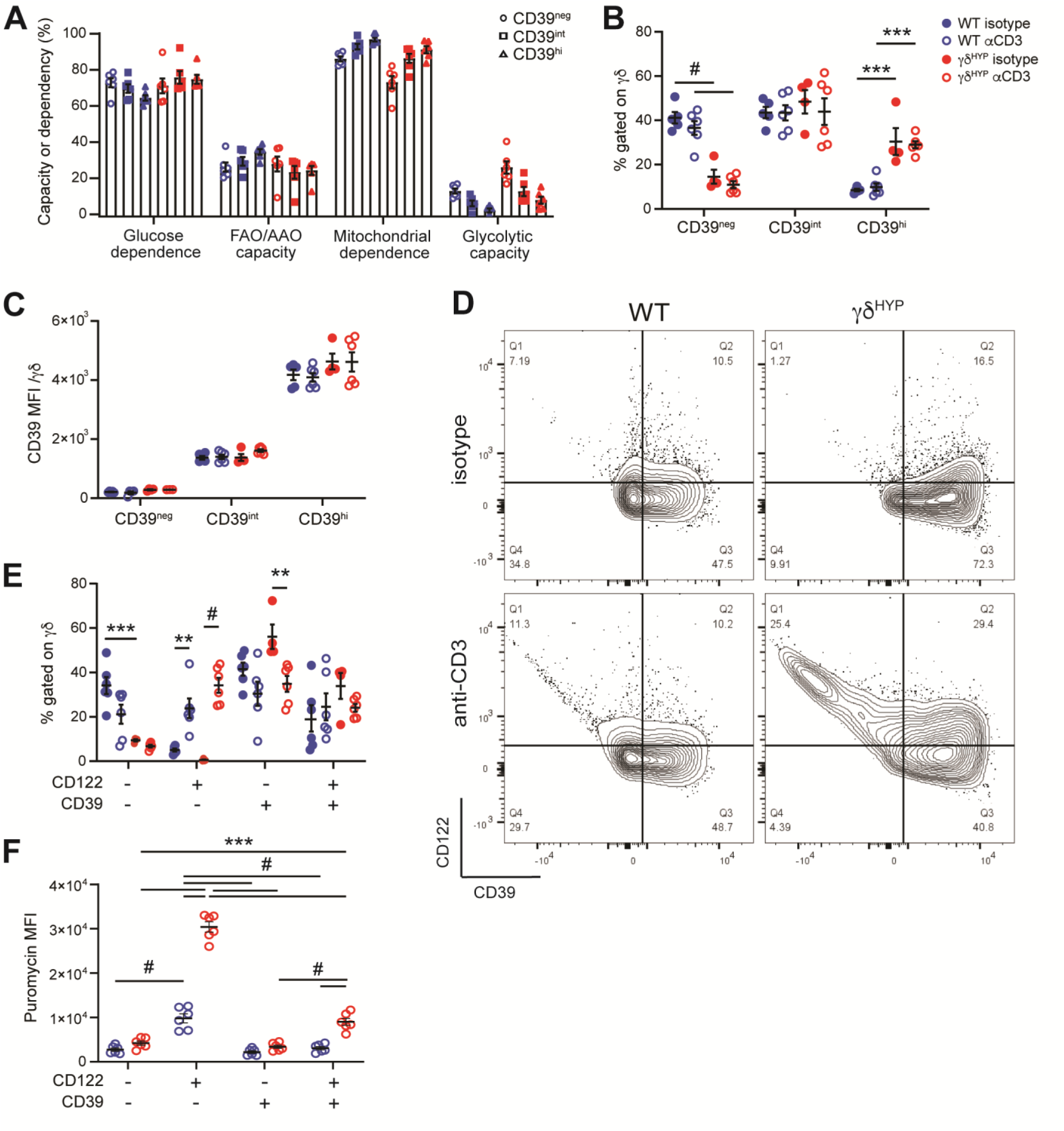
I*n vivo* activation induces the expansion of a CD39^neg^CD122^p05^γδ IEL population with increased protein production. WT and γδ^hvp^ mice were injected i.p with either isotype control or anti-CD3 for 16 h. (A) Percent capacity or dependency of different metabolic pathways among CD39 populations. (B) Frequency and (C) CD39 MFI of CD39^ne0^, CD39^int^, and CD39^hi^ yS IELs from WT and γδ^HYP^ mice. (D) Representative flow plots showing CD122 and CD39 expression following treatment. (E) Frequency and (F) puromycin MFI among CD122 and/or CD39-expressing y8 IELs. All data shown as mean ± SEM from at least 2 independent experiments. Each data point represents an individual mouse. n=6. Statistical analysis: (A-C,E,F) Two-way ANOVA with Tukey’s post hoc test. **P<0.01, ***P<0.001, #P<0.0001.

**Supplementary Table 1:**
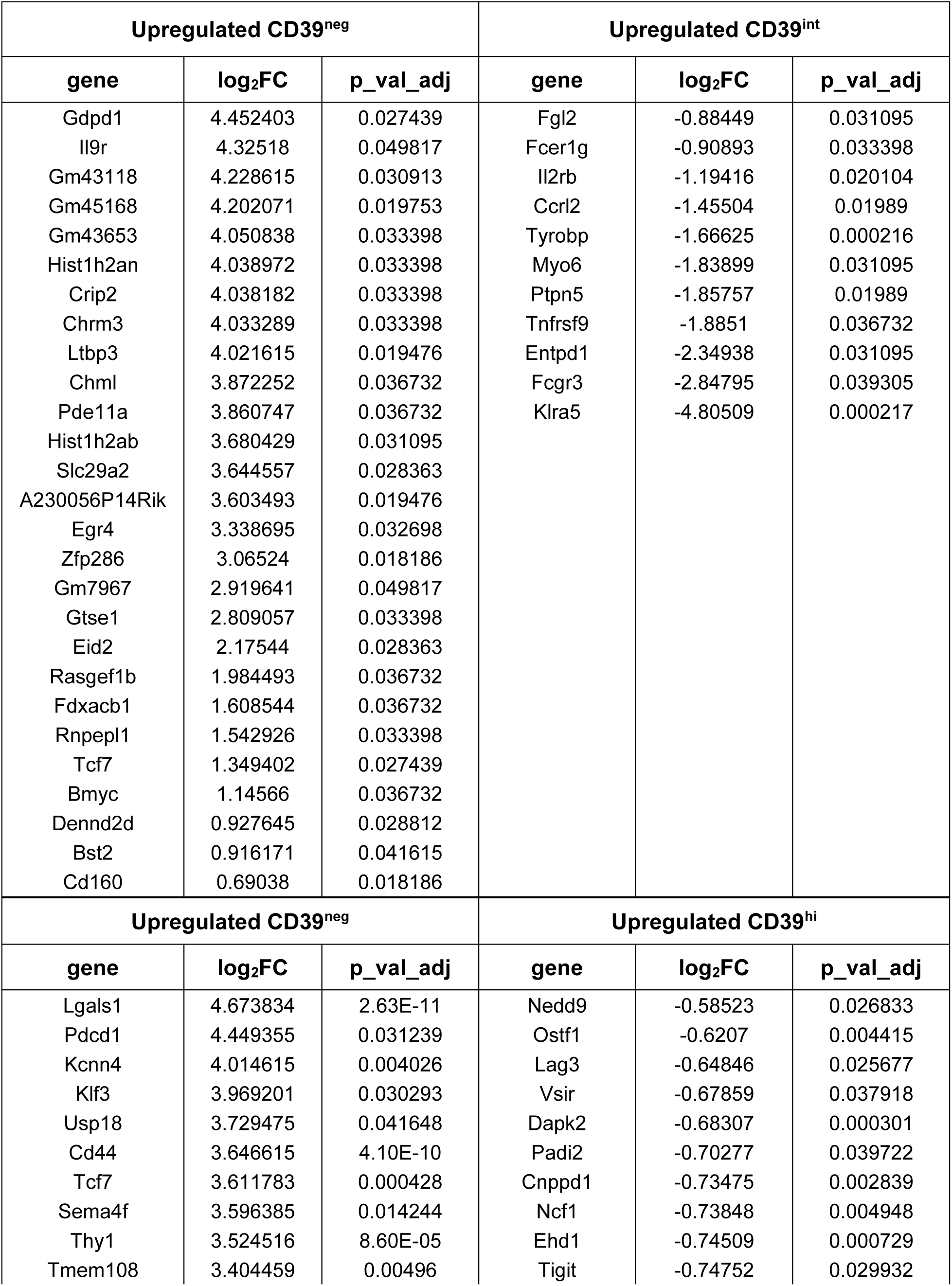
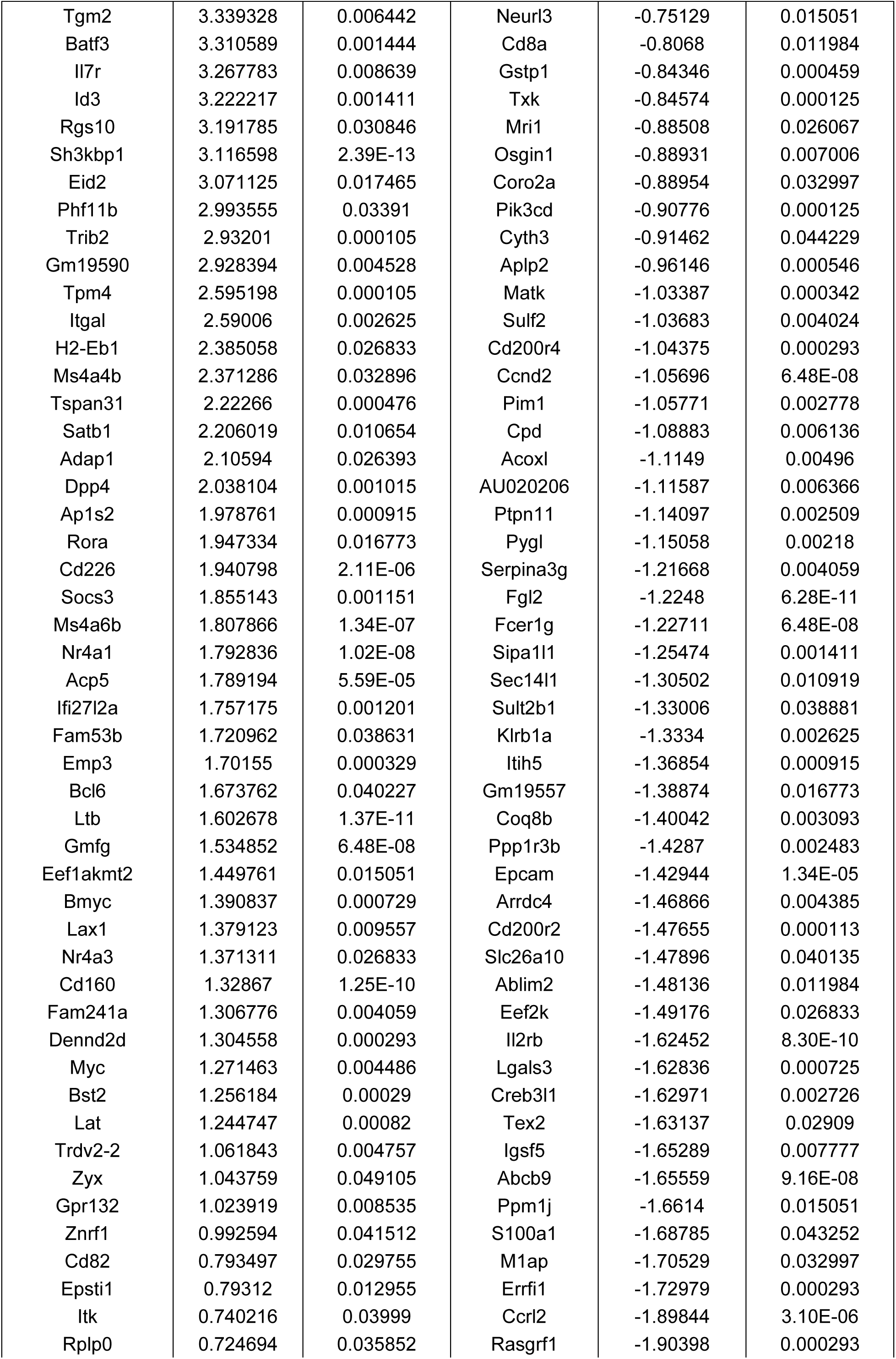
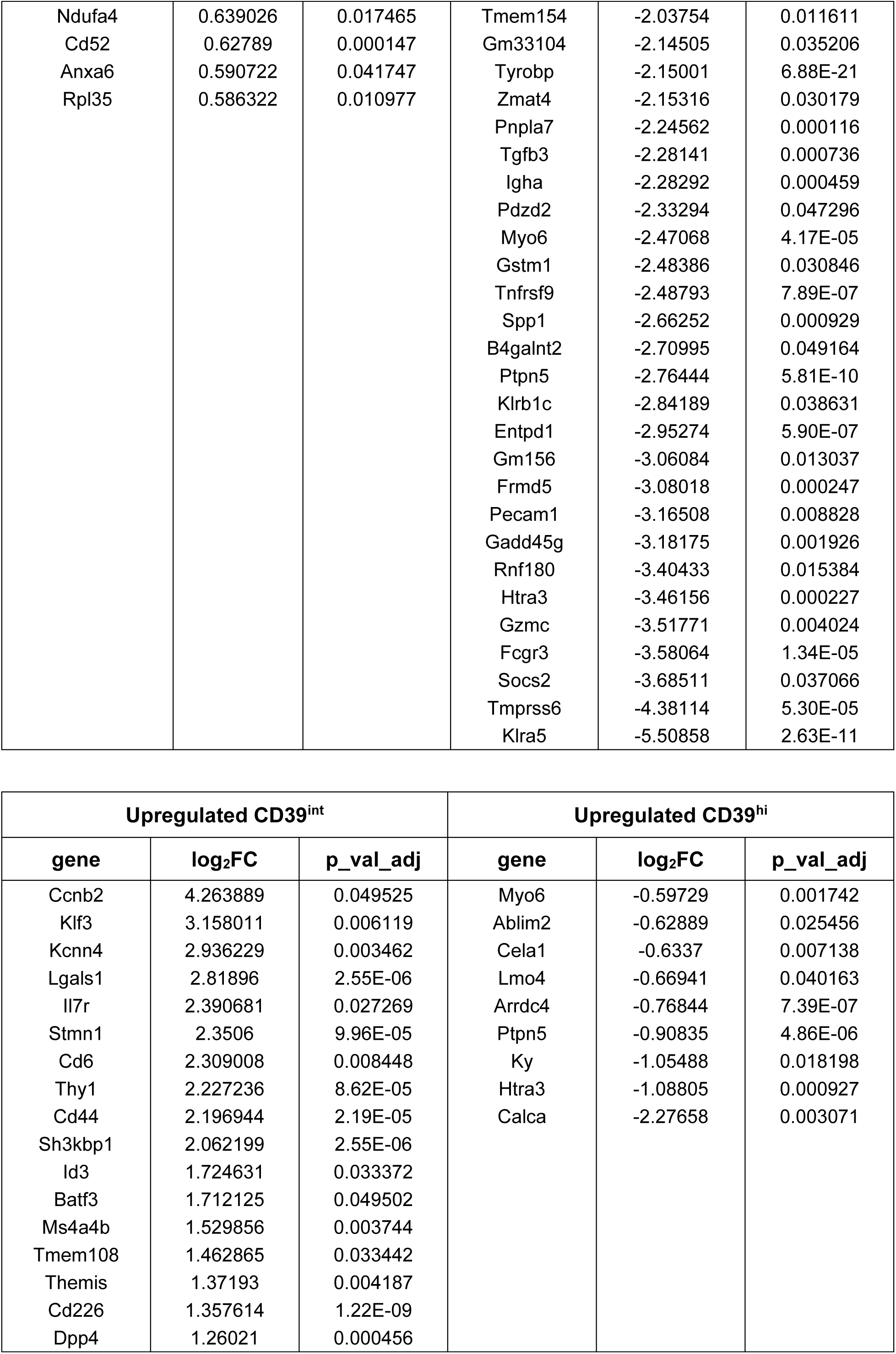
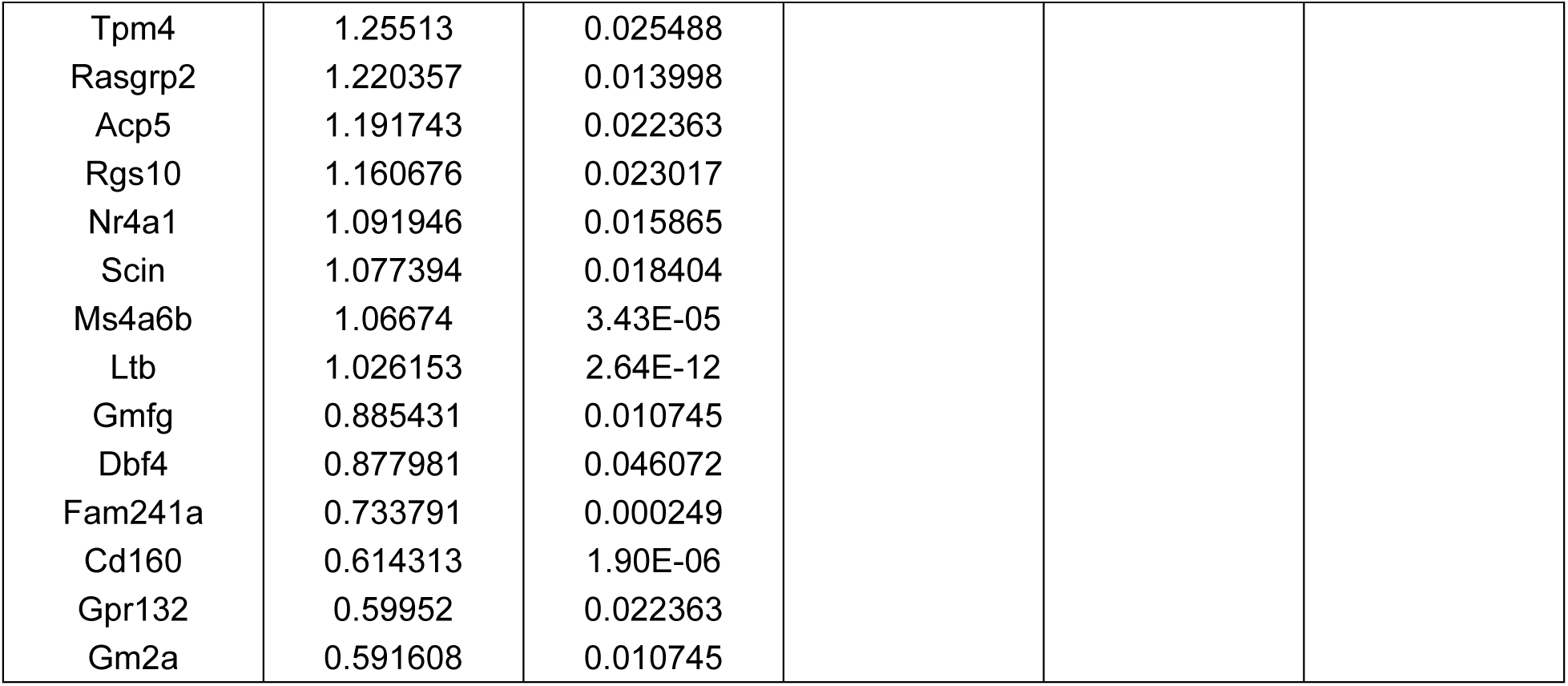
Pseudobulk analysis of DEGs between CD39-expressing. γδ **IEL subsets.** CITE-seq was performed on sorted CD3^+^ IELs from WT and γδ^HYP^ mice. CD39^neg^, CD39^int^, and CD39^hi^ γδ IELs were combined from WT and γδ^HYP^ mice and pseudobulk analysis was performed. Log_2_ fold change (FC) and adjusted p-values (p_val_adj) represented. Statistical analysis: ± 1.5-fold change, adjusted p-value of 0.05.

**Supplemental Table 2:**
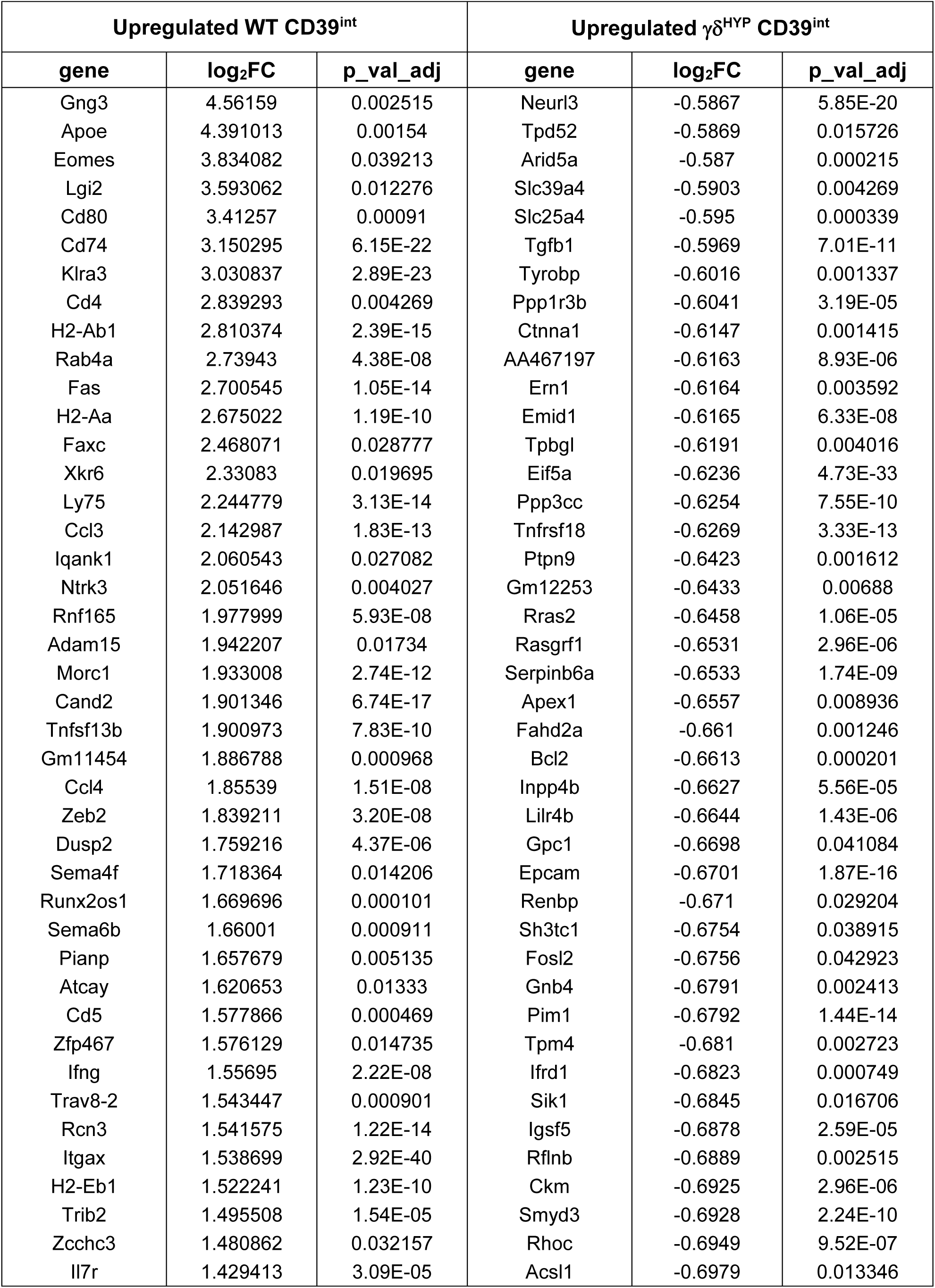
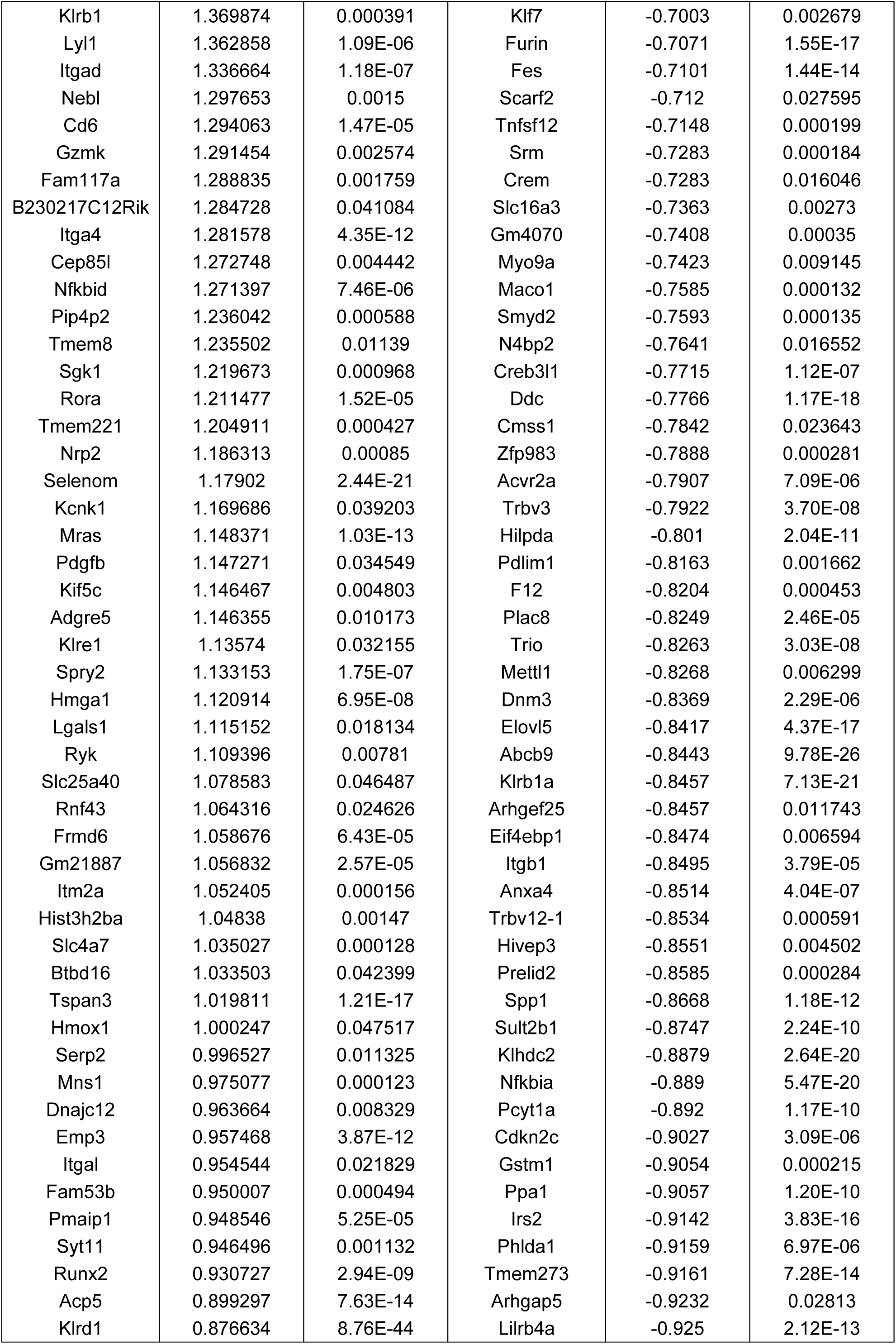
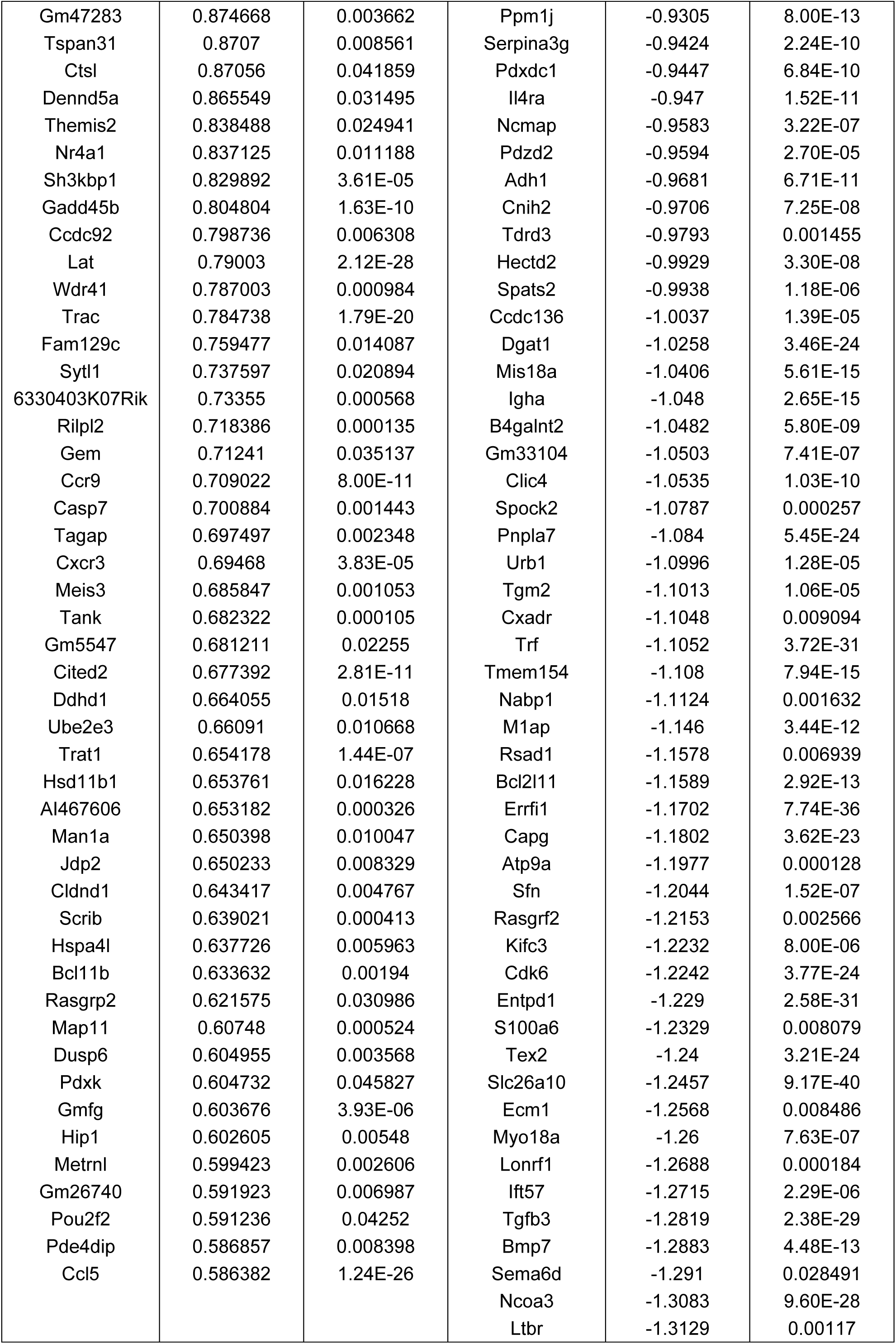
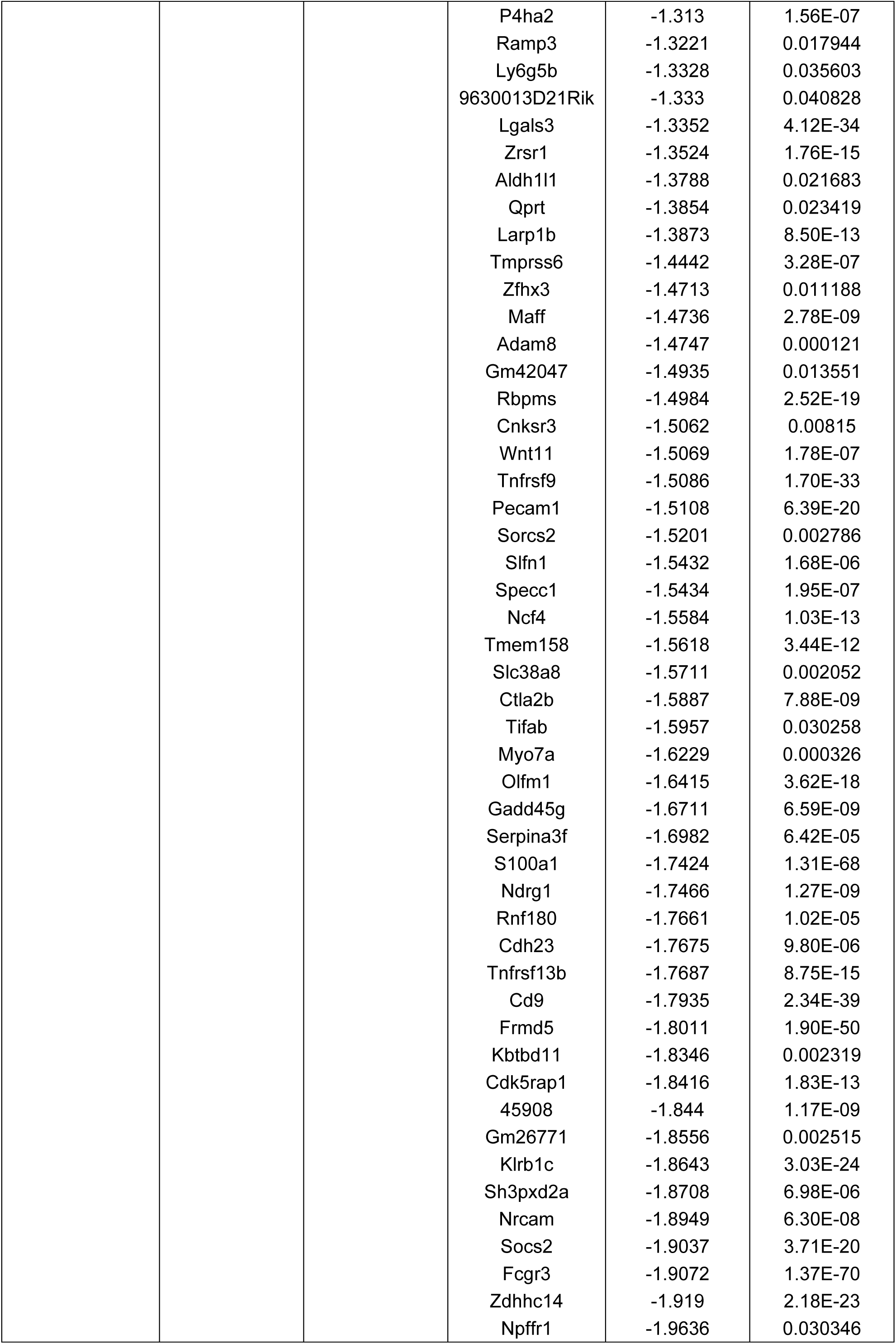
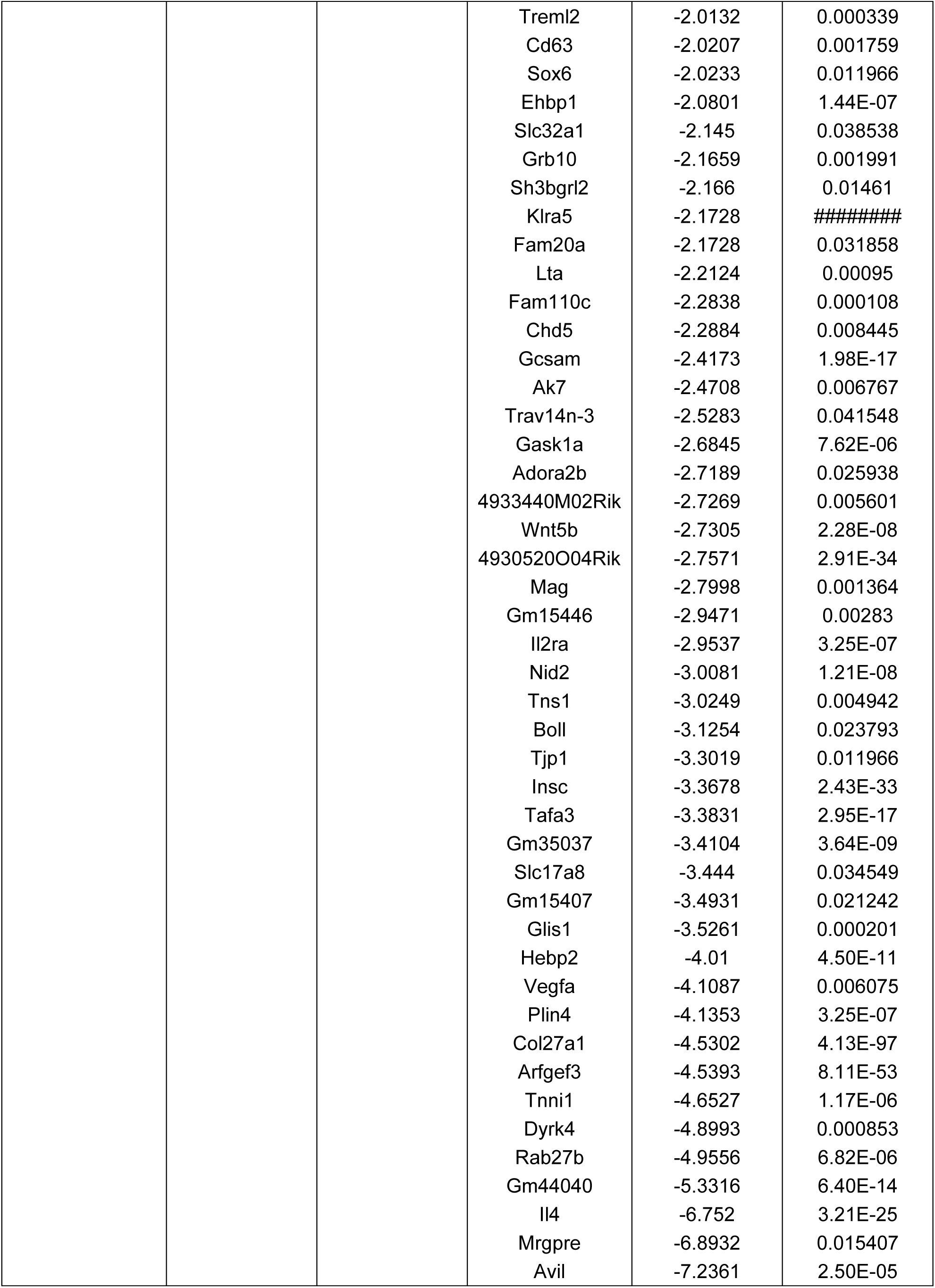
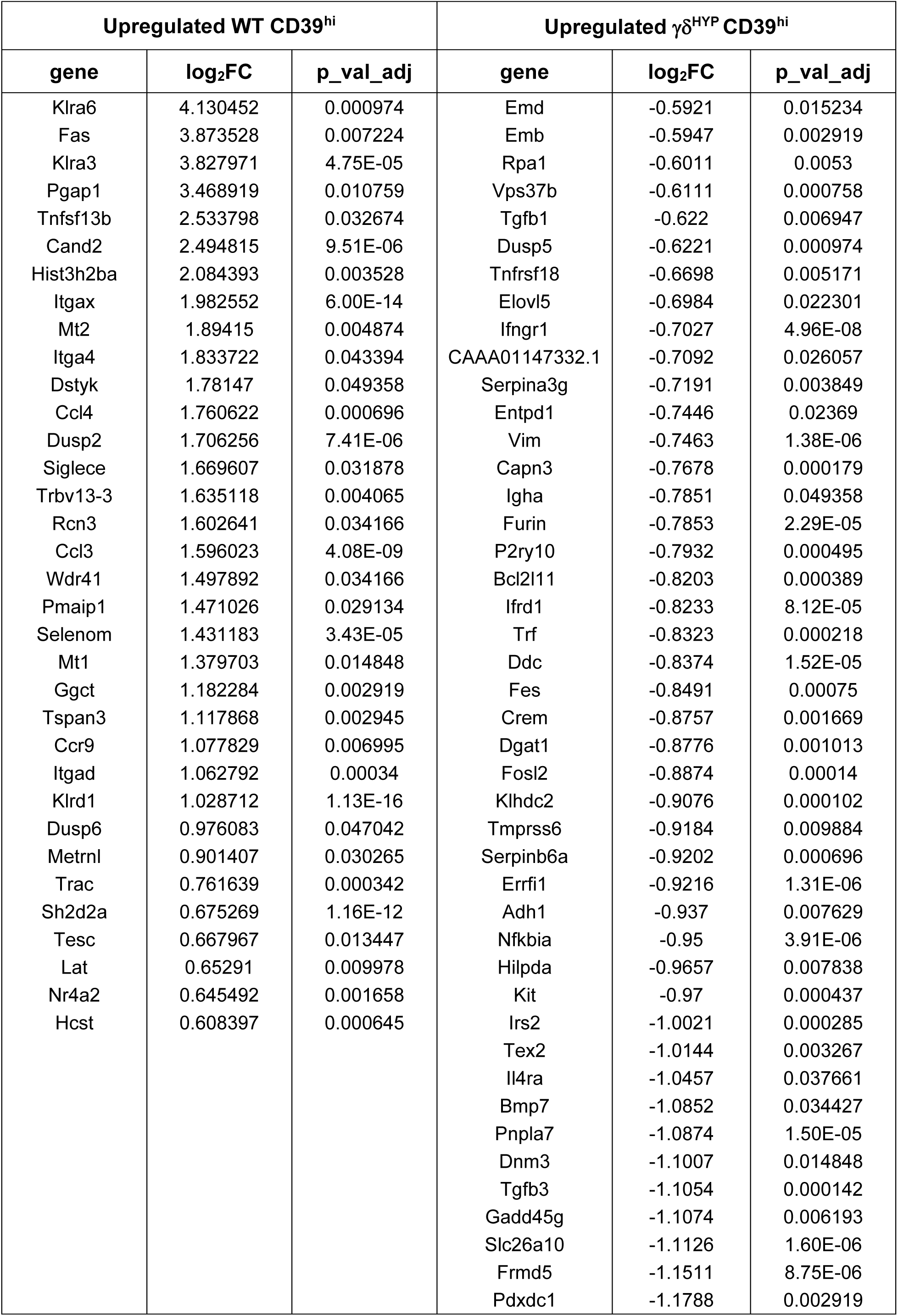
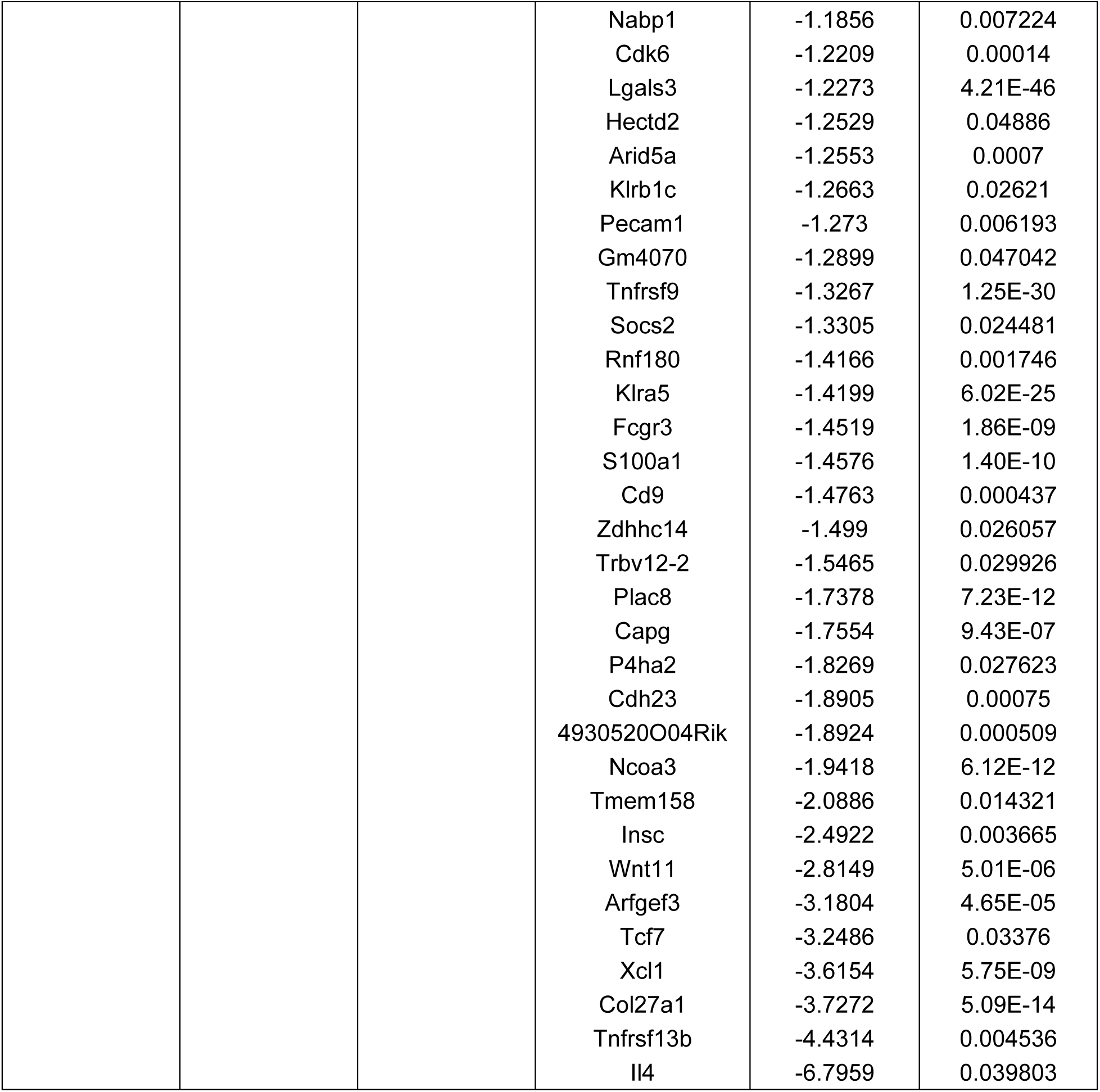
DEGs between phenotypic CD39^int^ and CD39^hi^. γδ **IELs.** Differential gene expression was assessed between CD39^int^ and CD39^hi^ γδ IELs from WT and γδ^HYP^ mice. Log_2_FC and p_val_adj represented. Statistical analysis: ± 1.5-fold change, adjusted p-value of 0.05.

